# Allosteric Regulation of RNA Affinity by Motif V-VI Coupling in West Nile Virus NS3 Helicase

**DOI:** 10.1101/2025.07.04.663198

**Authors:** Priti Roy, Martin McCullagh

## Abstract

The rise of flaviviral diseases, including West Nile virus (WNV), presents a growing threat to global public health and underscores the urgent need for new therapeutic strategies. The non-structural protein 3 helicase (NS3h) of the *Orthoflavivirus* genus, including WNV, is essential for viral replication and a promising antiviral target. Previously [Roy *et al.*, *Nucleic Acids Research*, 52, 13, 2024, 7447–7464], we showed that the motif VI loop (VIL) in WNV NS3h functions as a nucleotide valve, regulating ADP affinity during hydrolysis. In this study, we uncover an ATP-dependent coupling between nucleotide affinity at motif VIL and RNA affinity at motifs IVa and V, suggesting a coordinated mechanism of ssRNA translocation. Using microsecond-scale all-atom molecular dynamics simulations of hydrolysis-cycle intermediates, we find that key VIL residues (R461, R464) correlate strongly with RNA phosphate affinity of motif V. Structural analyses reveal an ATP-sensitive interaction between E413 (motif V) and R461 (motif VIL) that modulates the conformation of the motif V 3_10_-helix, thereby influencing RNA binding. This dynamic interaction is lost in catalytically deficient VIL mutants, which have been experimentally shown to impair hydrolysis and attenuate viral replication. These findings provide mechanistic insights into NS3h function and identify new opportunities for structure-based antiviral design.

## Introduction

West Nile Virus (WNV) remains a growing global public health concern. As an arthropod-borne (arbo) virus, WNV is maintained in a transmission cycle between vertebrate reservoir hosts and mosquito vectors. Human infection can result in a wide spectrum of outcomes, ranging from mild fever to severe neurological complications and, in some cases, kidney disease.^1–3^ WNV has caused multiple outbreaks^4,5^ and continues to expand into new geographic regions, with an increasing number of countries reporting cases.^6^ Despite significant advances in vaccine development over the past two decades,^7^ effective treatments for WNV infection remain elusive, and no antiviral drugs have been approved for clinical use.^7^ A promising approach to antiviral drug development focuses on targeting essential viral proteins^8,9^ which requires a comprehensive understanding of the structural mechanisms of viral protein functions.

The helicase domain of the non-structural protein 3 (NS3h) of WNV is one such target. WNV is a positive-sense single-stranded RNA (ssRNA) virus belonging to the the *Orthoflavivirus* genus. Following infection, the viral RNA is translated into viral proteins by the host cellular machinery.^10^ Viral replication proceeds through synthesis of a complementary negative-sense RNA strand, forming double-stranded RNA (dsRNA) intermediates. The separation of these intermediates, an essential step in replication, is driven by the helicase activity of NS3h. NS3h is highly conserved across *Orthoflavivirus* members such as DENV, ZIKV, and JEV, and mutations in this domain have been linked to increased virulence and adaptive evolution in WNV.^11^ NS3h unwinds dsRNA by translocating along the negative-sense strand in a 3^′^→5^′^ direction, powered by ATP hydrolysis. NS3h has therefore emerged as a high-value candidate for structure-based drug development.^12–15^ A detailed understanding of its structural mechanism is critical for the rational design of WNV-specific and broad-spectrum antivirals.

Functionally, NS3h coordinates RNA translocation and ATP hydrolysis through a structurally coupled mechanism. A DEAH-box helicase of superfamily 2 (SF2), NS3h is a monomeric protein comprising three domains (see Figure 1). Domains I and II, which adopt a Rossmann fold, form the active site for nucleoside triphosphatase (NTPase or ATPase) activity, commonly referred to as the ATP-pocket. Domain III, positioned opposite domains I and II, contributes to the formation of an RNA-binding cleft, or RNA-cleft. Although the ATP- and RNA-binding sites are spatially separated by ∼ 30 °A (center-of-mass distance), they exhibit strong functional interdependence: RNA binding enhances ATPase activity in the ATP-pocket,^16^ and ATP hydrolysis in turn drives conformational changes that mediate RNA translocation within the cleft.^17,18^ This reciprocal regulation suggests the existence of a structural mechanism that enables allosteric communication between the two sites.

**Figure 1:**
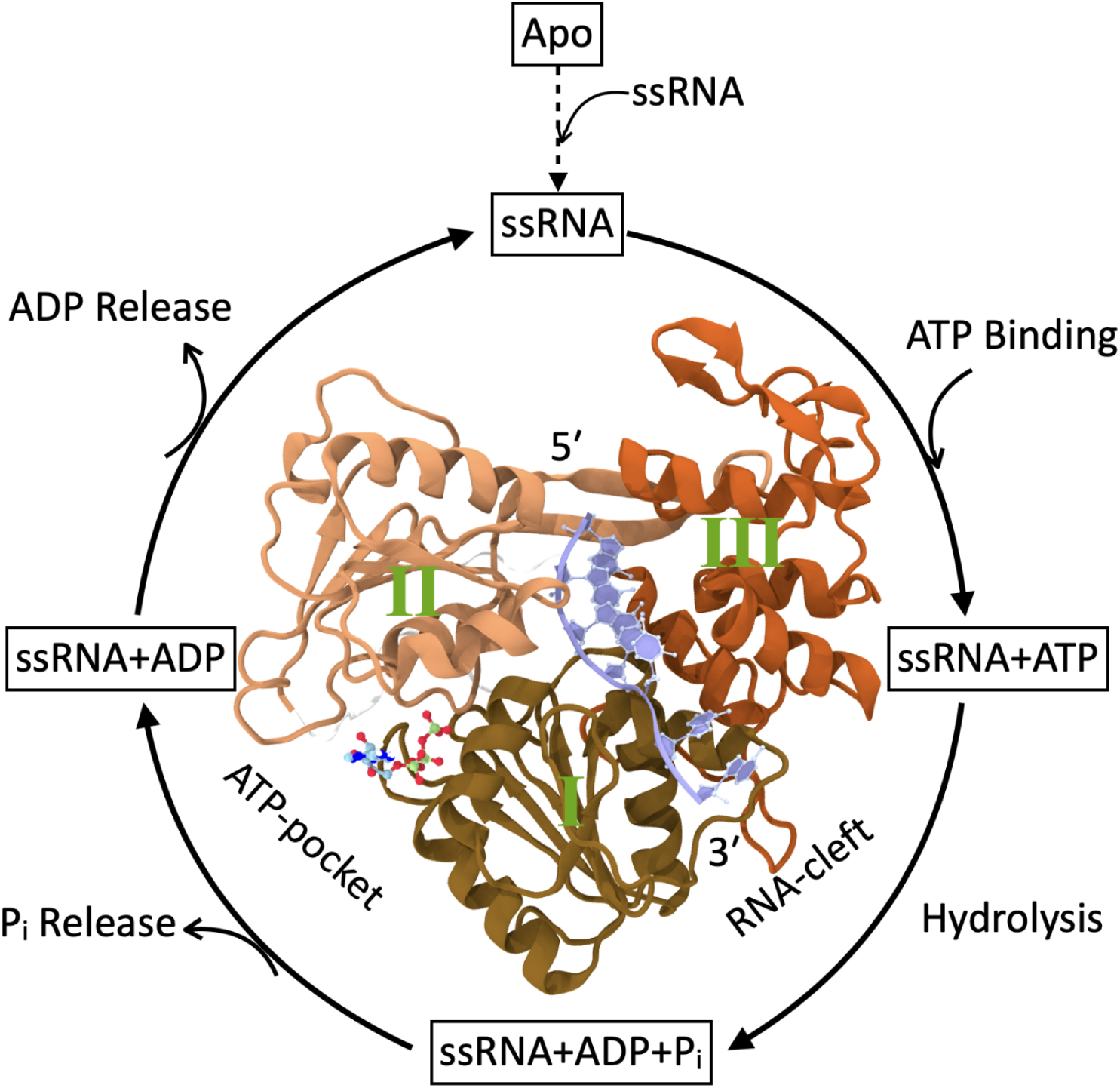
Schematic representation of the ATPase cycle of NS3 helicase (NS3h). Here we depict the intermediates states that NS3h transitions through during ATP hydrolysis cycle. The cycle starts when ssRNA enters RNA-cleft of the Apo NS3h and generates **ssRNA** state. After ssRNA binding, ATP enters the ATP-pocket and forms **ssRNA+ATP** state. Following hydrolysis, protein moves to **ssRNA+ADP+P**_i_ state. Next, P_i_ leaves the ATP-pocket and gives rise to **ssRNA+ADP** state. Finally, ADP releases and protein re-enter the **ssRNA** state to resume next cycle. Free energy released from this cycle drives translocation in 3^prime^ to 5^prime^ direction along the ssRNA to unwind dsRNA.

Mounting evidence suggests that the motif **VI** loop acts as a key regulatory element linking ATP hydrolysis to RNA translocation in flaviviral helicases. Our previous work combining molecular simulations and biochemical assays demonstrated that the motif **VI** loop in WNV NS3h modulates nucleotide binding enthalpy in a hydrolysis state-dependent manner, functioning as a nucleotide valve. ^19^ We also observed ATP-dependent changes in RNA affinity, a phenomenon reported in other members of the NS3 helicase family.^16,17,20,21^ Together, these observations support the hypothesis that ATP-dependent nucleotide affinity at motif **VI** loop may be coupled to changes in RNA affinity within the RNA-cleft.

This putative coupling likely involves motif **V**, which lies in close spatial proximity to motif **VI** loop and is structurally poised to mediate communication between the ATP-pocket and RNA-binding site. Motif **V** comprises a *β*-strand, a 3_10_-helix that interacts with ssRNA, and a coil facing the ATP-pocket. Prior studies have shown that ATP binding induces shifts in motif **V** toward the nucleotide-binding site in DENV NS3h,^22^ and that hydrogen bonding within its 3_10_-helix contributes to RNA unwinding and replication.^23^ Structural analyses of ZIKV NS3h revealed conformational variability in the coil region of motif **V**,^18^ and MD simulations suggest it samples distinct conformations across the hydrolysis cycle. Although our prior work found no direct enthalpic signature of hydrolysis state dependence in motif **V**, we hypothesize that it serves as a structural conduit, mediating ATP-dependent communication from motif **VI** to the RNA-cleft. The present study tests this hypothesis by analyzing ATP-dependent structural correlations between motifs **V** and **VI**, aiming to define a mechanistic link between ATP hydrolysis and RNA translocation, and to inform the development of NS3h-targeted antivirals.

## Methods

### Starting Structures and System Preparation

In this study, the D471E and D471L mutants, bound to ATP, ADP+P_i_, and ADP, were generated by modifying previously modeled wild-type WNV NS3h structures in complex with ssRNA.^19^ Specifically, the side-chain terminal atoms of D471 were replaced with those of glutamic acid (D471E) and leucine (D471L), respectively, while maintaining the overall protein architecture. The resulting mutant structures were solvated with TIP3P water molecules.^24^ To neutralize the system and maintain an ionic strength of 0.15 M, Na^+1^ and Cl^−1^ ions were added. A cubic box with a buffer distance of 20 °Afrom the protein surface was used. The dimensions of the resulting box were 107 °Aalong each axis. The simulations were conducted at neutral pH. For details on histidine protonation states, refer to the wild-type system preparation described in Roy et al., 2024.^19^

### Simulation Protocol

Mutant hydrolysis states were simulated similar to wild-type simulations^19^ with GPU-enabled AMBER18 software.^25^ In brief, each mutant state was minimized in the conjugate-gradient method, followed by heating at 310 K. After equilibration, the production run was performed for 5 *µ*s in the NPT ensemble. The integration time step was 2 fs and the coordinates were saved every 10 ps. Equation of motion was solved using ff14SB^26^ and ff99bsc0*χ*OL3^27,28^ as potential energy values for protein and ssRNA, respectively. The ATP,^29^ ADP,^29^ and Mg^2+ 30^ parameters were used, while the P_i_ (H_2_PO_4_^−^) parameter was adopted from a previous study by our group.^20^ In aggregates, we generated approximately 40 *µ*s of simulated frame and analyzed 60 *µ*s of simulated frame, including previously reported wild-type simulation data.^19^

### Analyses

#### Binding Enthalpy, E_inter_

Non-bonded linear interaction energy (E_inter_) between the protein and substrates i.e. the phosphate moieties of the ssRNA or ADP is computed from the NPT ensemble of each system which we denote as binding enthalpy. We computed electrostatic and van der Waals interactions as non-bonded interactions, using Cpptraj *lie* module with a long-range cutoff of 12 °A.

#### Pearson Correlation

The Pearson correlation measures the linear dependency of two variables. We computed the correlation coefficient which measures the extent of negative or positive correlation ranging from −1 to 1:

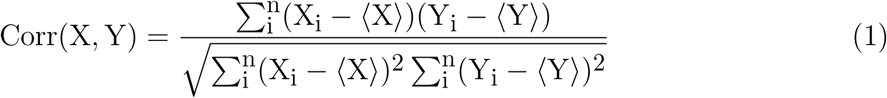

The ⟨X⟩ and ⟨Y⟩ denote the ensemble averages. Pearson’s correlation was evaluated for substrate binding affinities and Cartesian positions.

#### Hydrogen Bond (H-bond)

A hydrogen bond is considered when the two participating heavy atoms are electronegative and a hydrogen atom is attached to one of the heavy atom. The hydrogen bound electronegative atom acts as a donor, while the other atom acts as an acceptor. We used (i) a cutoff distance of 3 °Abetween the donor and acceptor atom, (ii) a cutoff distance of 1.1 °Abetweenthe donor and donor hydrogen atoms and (iii) an angle value higher than the 120^◦^ formed between the donor, hydrogen and acceptor atoms.

### Model Corroboration

Model hydrolysis states of the mutants are comparable to those of the corresponding wild-type states. Comparison of RNA contact sampling between mutants and wild-type shows that a large number of wild-type conserved contacts sample 70% - 100% in mutants (Supporting Information Table S1-S4). Similarly, we observe sampling of conserved ATP, ADP+P_i_ and ADP contacts, with a sampling probability higher than 70% for most of the wild-type contacts (Table S5-S7). However, a few conserved contacts are lost or samples below 20% in mutants. The octahedral coordination of Mg^2+^ is also maintained in the mutant hydrolysis states (Table S8). We computed the backbone dihedral angles of the wild-type and mutants and presented as a 2D histogram in the Supp. Info Figure S1. Mutations are incorporated by only changing the sidechain terminal moiety, and thus, these alterations would not have any influence on the secondary and tertiary structure. Similarly, the distributions of mutant states are comparable to those of wild-type states, largely sampling the *α*-helix region.

## Results and Discussion

The translocation activity of NS3h proceeds through a non-equilibrium, multistate hydrolysis cycle that couples RNA binding and ATP turnover. Crystal structures and biochemical studies have identified a sequence of conformational states corresponding to distinct steps in this ATPase cycle which we show in Figure 1. ^16,17,31,32^ The cycle begins when single-stranded RNA (ssRNA) binds within the RNA-cleft, while the ATP-binding site remains unoccupied; we refer to this state as ‘ssRNA’. RNA binding enhances nucleotide affinity,^33^ promoting ATP binding and formation of a pre-hydrolysis complex, ‘ssRNA+ATP’. ATP hydrolysis produces the post-hydrolysis-I state, ‘ssRNA+ADP+P_i_’, which then releases in-organic phosphate to yield the post-hydrolysis-II state, ‘ssRNA+ADP’. Finally, ADP release returns the protein to the initial ‘ssRNA’ state. These conformational transitions correspond to elementary steps of helicase motion, likely involving single-nucleotide translocation events.^32,34^ We adopt these state definitions for our analysis, which builds on prior work demonstrating ATP-dependent changes in nucleotide and RNA binding affinity. ^19^

In this context, we hypothesize that the motif **VIL** acts as a dynamic regulator that couples nucleotide affinity to RNA translocation during the hydrolysis cycle. This hypothesis is motivated by our previous study, which combined molecular dynamics simulations and biochemical assays to show that motif **VIL** plays a key role in controlling nucleotide entry, stabilization, and release.^19^ Although motif **VIL** is spatially distant from the ATP-binding site, mutation of a key residue impaired hydrolysis and viral replication, consistent with long-range allosteric regulation. We previously proposed that motif **VIL** acts as a nucleotide valve, modulating access to the ATP site across different hydrolysis states.^19^ Here, we investigate whether this valve-like behavior is structurally and functionally coupled to RNA affinity, thereby linking ATP turnover to translocation. Our analysis is organized into three parts: (1) the correlation between nucleotide (ADP) and RNA affinity; (2) residue-level interactions that bridge the ATP- and RNA-binding sites; and (3) structural coupling between motif **VIL** and RNA-binding motifs that varies across hydrolysis states.

### Correlation between nucleotide affinity at motif VI and RNA affinity

WNV NS3h contains eight conserved sequence motifs (**I**, **Ia**, **II**, **III**, **IV**, **IVa**, **V**, and **VI loop** [**VIL**]) that coordinate ATP hydrolysis and RNA translocation through substrate binding in two functionally distinct regions: the ATP-pocket and the RNA-cleft (Figure 2 A). Motifs **I**, **II**, and **III** form the nucleotide-binding pocket and catalyze ATP hydrolysis,^35–37^ while motifs **IV** and **IVa** reside in the RNA-cleft and participate in RNA unwinding and replication.^37,38^ Motif **V** is positioned between these regions and has been implicated in allosteric coupling.^18,37,38^ Motif **Ia**, which spans both regions, and motif **VIL**, the so-called arginine finger, are of particular interest: the former shows ATP-dependent motion in related helicases,^39,40^ and the latter stabilizes nucleotide binding and is essential for catalysis.^16,19^ Based on structure and prior functional studies, we classify motifs **I**, **II**, **III**, **V**, and **VIL** as ATP-pocket motifs and motifs **Ia**, **IV**, **IVa**, and **V** as RNA-cleft motifs. To assess whether ATP hydrolysis dynamically coordinates substrate binding in these regions, we investigated ATP-dependent coupling between nucleotide and RNA affinity across the hydrolysis cycle.

**Figure 2:**
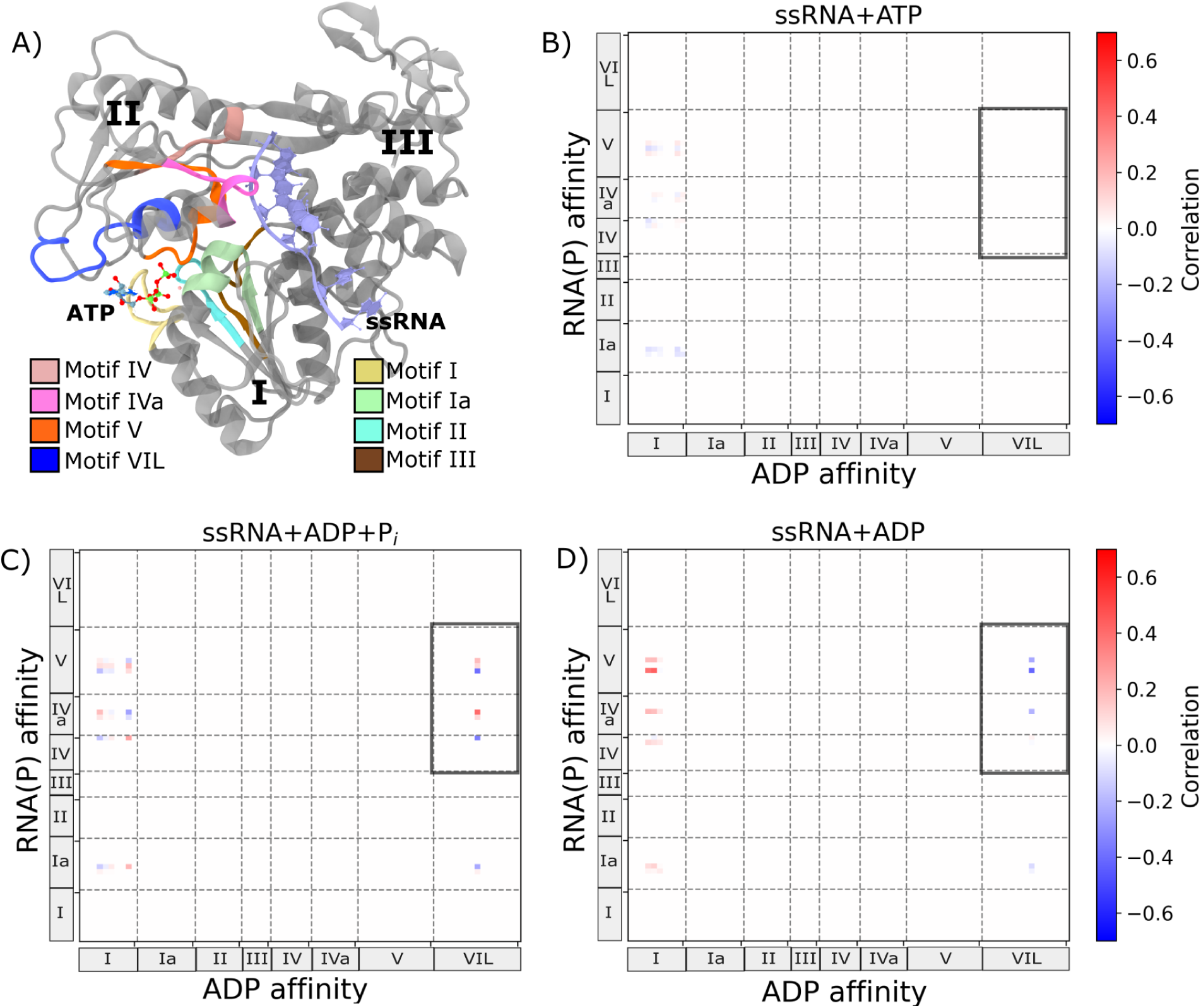
ATP-dependent correlation between nucleotide and RNA affinity in WNV NS3h. Binding affinity is defined as the interaction energy between a substrate and a motif residue within the protein. For nucleotides, we consider ADP (excluding the *γ*-phosphate group of ATP); for RNA, we quantify affinity based on interactions with phosphate moieties of single-stranded RNA (RNA(P)). (A) Structural model of WNV NS3 helicase (NS3h), highlighting locations of conserved motifs. NS3h contains eight conserved sequence motifs (I, Ia, II, III, IV, IVa, V, and VIL), all residing within domains I and II. (B–D) Residue-wise cross-correlation plots comparing nucleotide (ADP) and RNA(P) binding affinities for three hydrolysis states: (B) ssRNA+ATP (pre-hydrolysis), (C) ssRNA+ADP+P_i_ (post-hydrolysis I), and (D) ssRNA+ADP (post-hydrolysis II). Regions with marked changes in correlation across states are outlined with rectangular boxes.

Nucleotide and RNA binding affinities in NS3h become increasingly coupled as the hydrolysis cycle progresses, indicating that ATP hydrolysis drives coordinated substrate engagement across distant binding sites. Prior work has shown hydrolysis state-dependent nucleotide binding affinity in WNV NS3h, ^19^ as well as ATP-dependent changes in RNA affinity in both DENV4 and WNV helicases.^19,20^ These findings suggest that affinity for nucleotide and RNA may be coupled in a hydrolysis-state-dependent manner. To investigate this coupling, we computed residue-level cross-correlations between nucleotide and RNA binding affinities for all conserved motifs in three hydrolysis states: ssRNA+ATP, ssRNA+ADP+P_i_, and ssRNA+ADP (Figure 2B–D). RNA affinity was defined as the interaction energy between motif residues and the phosphate moieties of the RNA backbone (RNA(P)). To ensure consistency across states, nucleotide affinity was calculated using interaction energies with ADP, which is present in all three states. This analysis revealed that nucleotide and RNA affinities become increasingly correlated as the hydrolysis cycle progresses. In the ssRNA+ATP state, correlation coefficients ranged from –0.095 to 0.073 (Figure 2B), indicating weak coupling. In contrast, the correlation range broadened significantly in the ssRNA+ADP+P_i_ (–0.396 to 0.406; Figure 2C) and ssRNA+ADP (–0.417 to 0.433; Figure 2D) states. This ATP-dependent increase in coupling strength supports a model in which nucleotide binding at the ATP-pocket influences RNA affinity within the cleft, and vice versa.

Among ATP-pocket motifs, **I** and **VIL** show the strongest coupling to RNA affinity, but with distinct state-dependent profiles—motif **VIL**, in particular, exhibits behavior consistent with an allosteric regulatory role. Motif **I**, a core component of the ATP-binding pocket, shows increasing correlation with RNA(P) affinity as the cycle progresses. In the ss-RNA+ATP state, correlation coefficients between motif **I** and RNA(P) range from –0.095 to 0.073, indicating weak coupling (Figure 2B). This correlation becomes more pronounced in the ssRNA+ADP+P_i_ state (–0.254 to 0.252; Figure 2C), and shifts further toward positive correlation in the ssRNA+ADP state (–0.038 to 0.433; Figure 2D). In contrast, motifs **II**, **III**, and **V** show no significant correlation with RNA affinity in any hydrolysis state, suggesting they primarily support hydrolysis rather than substrate coupling. Motif **VIL** displays a distinct pattern: no correlation in the ssRNA+ATP state (Figure 2B), but strong bidirectional correlations emerge in the ssRNA+ADP+P_i_ state (–0.396 to 0.406; Figure 2C), which then shift toward predominantly negative correlations in the ssRNA+ADP state (–0.417 to 0.036; (Figure 2D)). This ATP-dependent behavior of motif **VIL** supports its proposed role as a nucleotide valve that dynamically couples nucleotide affinity to RNA binding through the course of the hydrolysis cycle.

ATP-dependent coupling between nucleotide and RNA affinity is also evident in the RNA-cleft motifs, particularly motifs **IV**, **IVa**, and **V**. These motifs show increasing correlation with the nucleotide affinity of motif **I** and motif **VIL** as the hydrolysis cycle progresses. In the ssRNA+ATP state (Figure 2B), correlations between nucleotide affinity at motif **I** and RNA(P) affinity are generally weak across RNA-cleft motifs (e.g., –0.093 to 0.0 for **Ia**, –0.080 to 0.033 for **IV**), and no notable correlation is observed for motif **VIL**. In the ssRNA+ADP+P_i_ state (Figure 2C), stronger correlations emerge. RNA affinity of motifs **IV** and **IVa** show the highest correlation with motif **I** nucleotide affinity (ranging from –0.254 to 0.252), while motif **V** also shows moderate coupling. Strikingly, motif **VIL** shows emergent bidirectional correlations with RNA affinity in motifs **IV**, **IVa**, and **V**, including a peak correlation of 0.406 with motif **IVa**. By the ssRNA+ADP state (Figure 2D), the coupling landscape diverges, revealing distinct roles for motifs **I** and **VIL**. Correlation between motif **I** and motif **V** reaches a maximum (up to 0.433), whereas correlations with motif **VIL** become predominantly negative, particularly for motifs **IVa** and **V**. These trends highlight a reorganization of coupling between nucleotide- and RNA-binding regions during the hydrolysis cycle and underscore the potential role of motif **V** as a key mediator in this allosteric communication.

Motif **VIL** exhibits more pronounced and state-specific coupling to RNA affinity than motif **I**, suggesting a unique mechanism for linking ATP hydrolysis to RNA engagement. From the ssRNA+ATP to the ssRNA+ADP+P_i_ state, the correlation between motif **VIL**’s ADP affinity and RNA(P) affinity increases sharply—particularly with RNA-cleft motifs **IV**, **IVa**, and **V**—exceeding the magnitude of change observed for motif **I**. As the system transitions to the ssRNA+ADP state, motif **I**’s correlation with RNA affinity diminishes for most RNA-cleft motifs, persisting only for motif **V**. In contrast, motif **VIL** maintains a strong correlation with motif **V** and shows moderate coupling with motif **IVa**, while correlations with motif **IV** is lost. These state-specific patterns are consistent with our previous findings that highlighted motif **VIL**’s central role in modulating nucleotide affinity across the hydrolysis cycle. ^19^ Taken together, these suggest that motif **VIL** may function as a conduit for transferring free energy from ATP hydrolysis to RNA translocation via selective coupling with RNA-binding elements.

Disruption of the ATP-dependent correlation between motif **VIL** and RNA affinity impairs helicase function in WNV NS3h. Molecular dynamics simulations of hydrolysis-state models for two motif **VIL** mutants—D471E (Figure S2) and D471L (Figure S3)—reveal a loss of the strong ADP–RNA affinity correlation observed in the wild-type enzyme. These mutants, previously shown to exhibit severely reduced nucleotide binding affinity,^19^ fail to maintain the coordinated interaction between ADP binding and RNA affinity. Notably, the correlation between motif **I** ADP affinity and RNA(P) affinity varied across the mutants, in some cases strengthening and in others weakening relative to wild-type. However, this variability did not compensate for the loss of motif **VIL** function, suggesting that the ATP-dependent coupling mediated specifically by motif **VIL** is critical for the enzymatic activity. Together, these findings establish motif **VIL** as a critical structural element for coupling ATP hydrolysis to RNA translocation in WNV NS3h.

### Correlation between nucleotide affinity at motif VI and RNA affinity at motif V

ATP-dependent changes in phosphate-specific RNA affinity at motifs **IVa** and **V** identify key contacts that may coordinate with nucleotide binding at motif **VIL** during translocation. In particular, motifs **IVa** and **V** exhibit notable changes in affinity for the first three RNA phosphates (P_1_–P_3_) across the hydrolysis cycle. To identify the most functionally relevant RNA-binding sites, we calculated changes in interaction energy (ΔE_inter_) between each RNA-cleft motif and individual phosphate groups across successive states. Figure S4 shows phosphate-specific ΔE_inter_ values for motifs **Ia**, **IV**, **IVa**, and **V**, revealing significant shifts in affinity at P_1_ and P_3_ for motif **IVa**, and at P_2_ for motif **V**. Figure 3 presents residue-level phosphate affinity changes over the course of the hydrolysis cycle: each row (from top to bottom) shows transitions corresponding to ATP binding (A–C), hydrolysis (D–F), P_i_ release (G–I), and ADP release (J–L). These data identify T409 and D410 in motif **V** as key contributors to changes in P_2_ affinity, while R388 and K389 in motif **IVa** drive affinity shifts at P_1_ and P_3_. Representative structures in Figure 4 illustrate the evolving positions of these residues and phosphates across hydrolysis states, providing a structural basis for ATP-dependent coupling between RNA- and nucleotide-binding sites.

**Figure 3:**
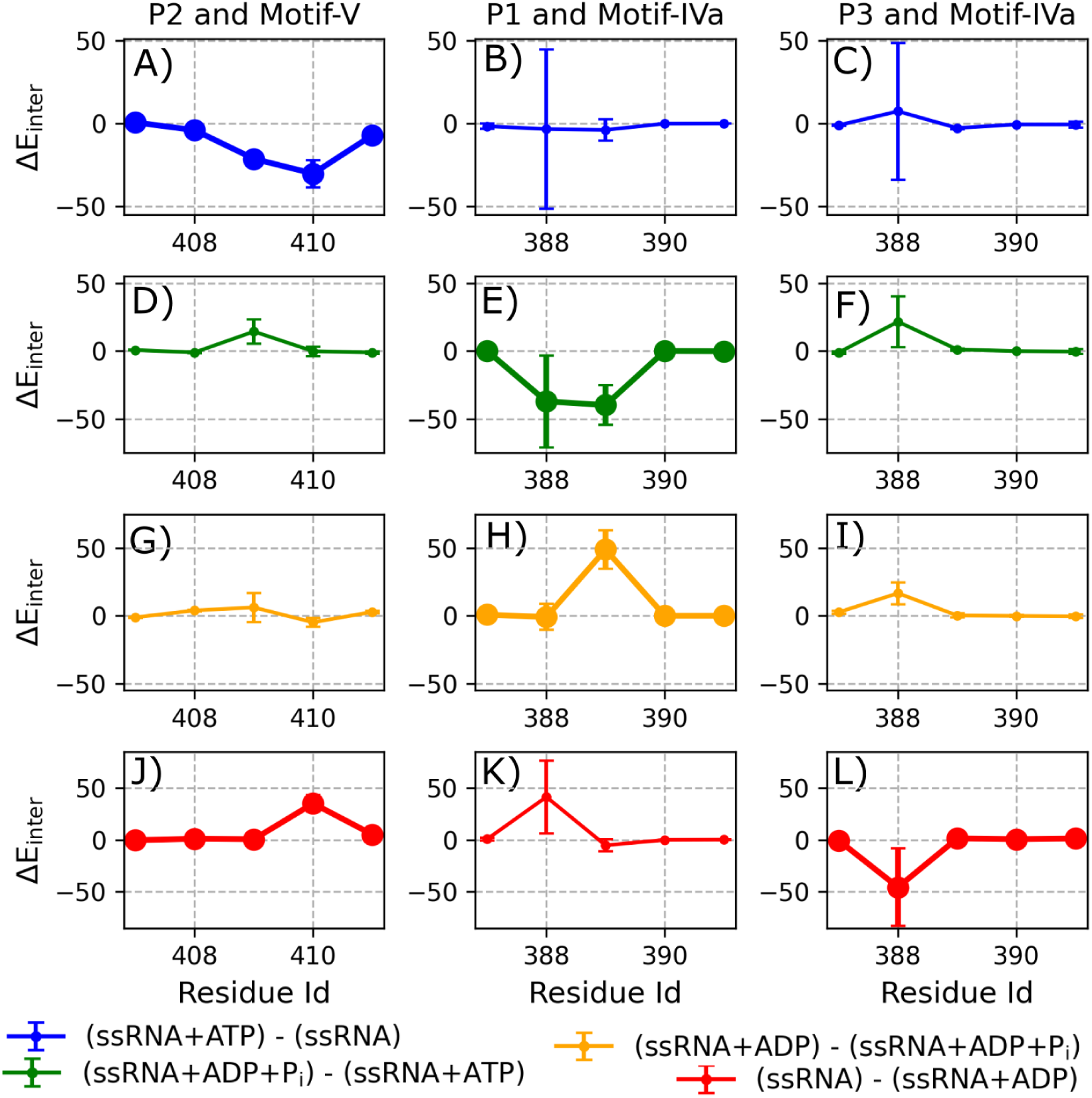
ATP-dependent changes in RNA phosphate (RNA(P)) affinity for motifs IVa and V. Each panel shows the residue-wise change in RNA phosphate affinity (ΔE_inter_, in kcal·mol^−1^) for key contacts between protein motifs and ssRNA phosphates during transitions between hydrolysis states. Left, middle, and right columns correspond to changes in affinity between motif **V** and P_2_, motif **IVa** and P_1_, and motif **IVa** and P_3_, respectively. Values of ΔE_inter_ reflect the difference in ensemble-averaged interaction energies computed over full NPT trajectories for each state. Error bars represent the propagated standard deviation of the difference between two ensemble means, where each mean is computed from 50,000-frame chunks. (A–C) Transition from ssRNA to ssRNA+ATP. (D–F) Transition from ssRNA+ATP to ssRNA+ADP+P_i_. (G–I) Transition from ssRNA+ADP+P_i_ to ssRNA+ADP. (J–L) Transition from ssRNA+ADP (previous cycle) to ssRNA (next cycle). Thicker lines indicate residues exhibiting significant affinity changes during a given transition. See *Analysis* section for computational details.

**Figure 4:**
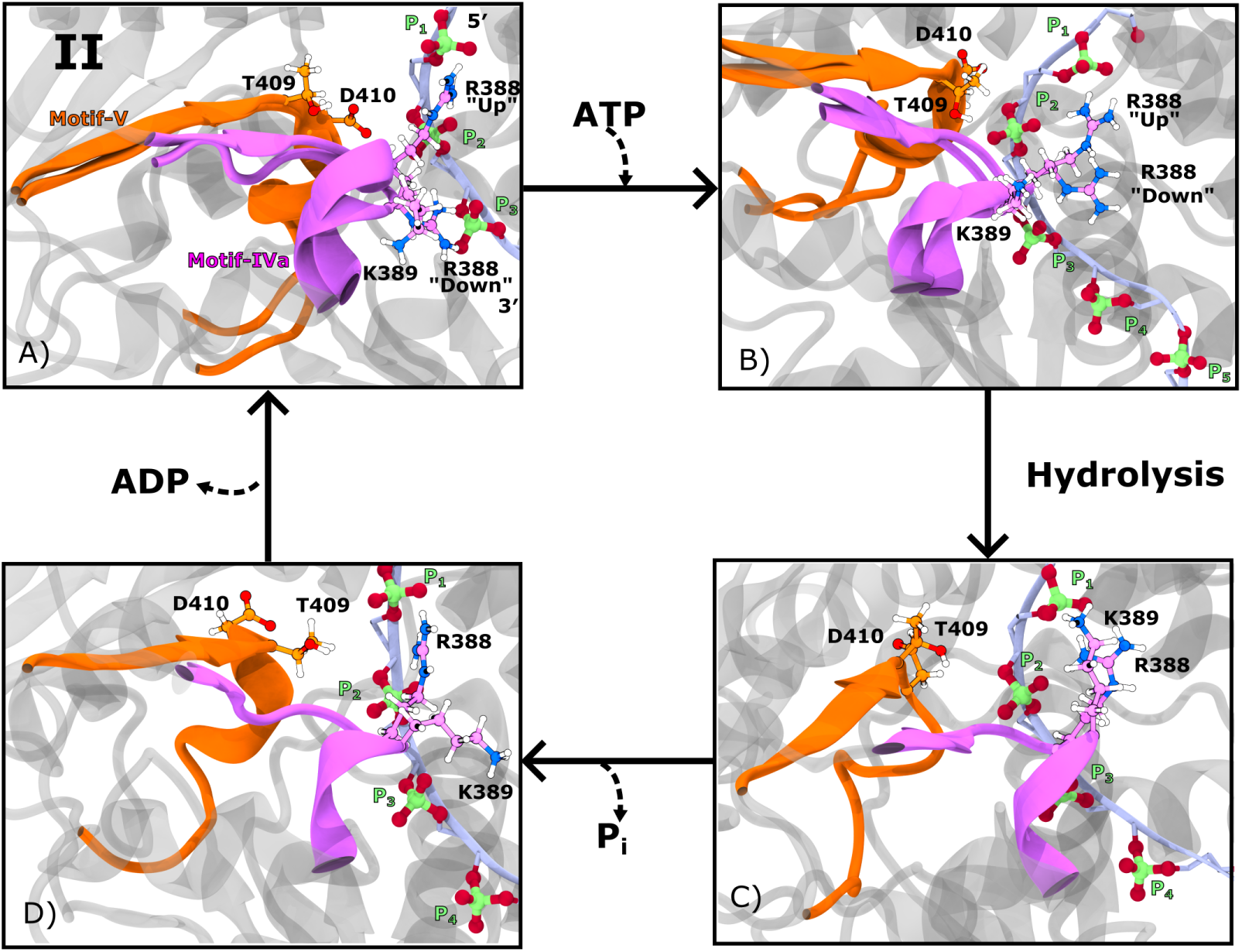
Representative structures of the four hydrolysis states illustrating motif IVa and V interactions with RNA phosphates. Key residues from motif **IVa** (R388 and K389) and motif **V** (T409 and D410) are shown, as these exhibit significant changes in RNA phosphate affinity (P_1_, P_2_, and P_3_) across the hydrolysis cycle. Motif **IVa** and its residues are colored orange; motif **V** and its residues are colored magenta. The remainder of the protein is shown in translucent grey. The ssRNA is depicted as an ice-blue ribbon, with its phosphate moieties shown in CPK representation (oxygen: red, phosphorus: green). (A) Representative structure of the ssRNA state, with two overlaid snapshots demonstrating R388 sampling both the ‘up’ conformation (near P_1_) and the ‘down’ conformation (near P_3_). D410 is positioned near P_2_, while K389 and T409 remain distant from RNA phosphates. (B) Structure of the ssRNA+ATP state showing post-ATP binding rearrangements. R388 continues to sample both conformations, while D410 detaches from P_2_ and T409 moves closer to occupy its position. (C) Structure of the ssRNA+ADP+P_i_ state following hydrolysis. R388 transitions to favor binding P_1_, and K389 also forms a close interaction with P_1_. Both motif **V** residues are detached from the RNA. (D) Structure of the ssRNA+ADP state following P_i_ release. K389 detaches from P_1_, while R388 maintains the ‘up’ conformation.

ATP binding induces asymmetric conformational responses in motifs **V** and **IVa**, revealing distinct mechanisms of phosphate engagement. Upon ATP binding, motif **V** forms stable contacts with phosphate P_2_, whereas motif **IVa** continues to fluctuate between interactions with P_1_ and P_3_. Specifically, motif **V** residue T409 exhibits a calculated ΔE_inter_ of −21.25 ± 1.42 kcal·mol^−1^ with P_2_, while D410 shows a stronger interaction of −30.16 ± 8.17 kcal·mol^−1^ (Figure 3A). Notably, this apparent increase in affinity for D410 reflects a reduction in repulsive interactions: its interaction energy shifts from 22.24±3.70 kcal·mol^−1^ in the ssRNA state to 7.83 ± 1.91 kcal·mol^−1^ in the ssRNA+ATP state. In contrast, motif **IVa** does not exhibit a net change in affinity for P_1_ (Figure 3B), although large fluctuations are observed in R388’s interaction with both P_1_ (ΔE_inter_ = −3.36 ± 47.86 kcal·mol^−1^) and P_3_ (ΔE_inter_ = 7.34 ± 41.10 kcal·mol^−1^) (Figure 3C). These findings suggest that ATP binding stabilizes motif **V** near P_2_, while R388 in motif **IVa** dynamically samples both an ‘up’ conformation near P_1_ and a ‘down’ conformation near P_3_ (Figure 4B).

Following ATP hydrolysis, the RNA-cleft reorganizes: motif **V** detaches from phosphate P_2_, while motif **IVa** transitions into a stably bound conformation at P_1_. In this ssRNA+ADP+P_i_ state, we observed a notable weakening in motif **V**–P_2_ interactions, with T409 showing a ΔE_inter_ of 14.38±8.90 kcal·mol^−1^ (Figure 3D), indicative of reduced affinity. D410 remains distant from P_2_ and exhibits no significant change, consistent with sustained low repulsion. In contrast, motif **IVa** forms stronger and more consistent interactions with P_1_: R388 and K389 exhibit ΔE_inter_ values of −37.23 ± 33.81 and −39.90 ± 14.57 kcal·mol^−1^, respectively (Figure 3E-F). The large uncertainty for R388 reflects its prior fluctuation in the ATP-bound state. Additionally, we observe a weakening of R388–P_3_ affinity (ΔE_inter_ = 21.62 ± 18.82 kcal·mol^−1^), indicating a shift away from P_3_ and toward stable binding at P_1_. Together, these changes define a hydrolysis-dependent structural reorganization, with motif **V** detached from P_2_ and motif **IVa** adopting an ‘up’ conformation stabilized at P_1_ (Figure 4C). A similar hydrolysis-induced R388 ‘up’ conformation was reported in simulations of DENV4 NS3 helicase,^20^ although in that system the ‘up’ conformation was also favored in the ATP-bound state—an important contrast to our findings in WNV NS3h.

Following P_i_ release, motif **IVa** begins to disengage from phosphate P_1_, while motif **V** remains detached from P_2_. During the transition from the ssRNA+ADP+P_i_ state to the ssRNA+ADP state, we observed no significant change in P_2_ affinity for either T409 or D410 of motif **V** (Figure 3G), suggesting that T409 maintains a weak interaction and D410 remains non-repulsive. For motif **IVa**, R388 shows no net change in affinity for P_1_, consistent with a continued strong interaction (Figure 3H), and remains weakly associated with P_3_ (ΔE_inter_ = 16.66 ±8.04 kcal·mol^−1^; Figure 3I). In contrast, K389 shows a notable weakening in affinity for P_1_, without a compensatory increase in affinity for P_3_, indicating a net loss of RNA interaction. The smaller error bars in Figures 3H and 3I, relative to those in the earlier state (Figures 3E–F), suggest reduced fluctuations in the ssRNA+ADP+P_i_ state. Taken together, these changes yield a partially detached motif **IVa**, in which R388 retains its ‘up’ conformation at P_1_, while K389 becomes RNA-free. Motif **V** remains dissociated from P_2_ throughout this state (Figure 4D).

Following ADP release, motif **V** re-engages partially with P_2_, while motif **IVa** resumes interactions with both P_1_ and P_3_. During the transition from the ssRNA+ADP state to the ssRNA state, we observed a significant change in the interaction energy between D410 and P_2_ (ΔE_inter_ = 35.20 ± 8.05 kcal·mol^−1^) (Figure 3J), while no net change was detected for other motif **V** residues. This shift reflects a weakening of D410’s interaction with P_2_, driven by an increase in repulsive affinity in the ssRNA state (22.24 ± 3.70 kcal·mol^−1^) relative to the ssRNA+ADP state (10.20 ± 2.73 kcal·mol^−1^). This repulsion results from D410 reattaching to P_2_ while T409 detaches (Figure 4A). For motif **IVa**, R388 exhibits substantial changes in phosphate affinity: a weakening of interaction with P_1_ (ΔE_inter_ = 41.52 ± 35.21 kcal·mol^−1^) and a strengthening with P_3_ (ΔE_inter_ = −45.62 ± 37.41 kcal·mol^−1^; Figure 3K-L). The large error bars associated with R388 reflect continued fluctuation between P_1_ and P_3_, indicating that this residue resumes sampling of both the ‘up’ and ‘down’ conformations in the ssRNA state (Figure 4A). Together, these state-specific conformational changes provide the structural basis for directional phosphate engagement during translocation.

The ATP-dependent RNA phosphate affinity and structural shifts of motifs **IVa** and **V** support an inchworm-style mechanism for RNA translocation during the hydrolysis cycle. Considering the 3^′^ to 5^′^ translocation direction characteristic of SF2 helicases, we propose the following model. The cycle begins with ATP binding, during which motif **IVa** releases phosphate P_n_ (corresponding to P_3_) and advances toward P_n+2_ (P_1_), while motif **V** engages P_n+1_ (P_2_). After hydrolysis, motif **IVa** binds to P_n+2_, and motif **V** detaches from P_n+1_. Upon P_i_ release, motif **IVa** prepares to release P_n+2_, and motif **V** remains unbound. Following ADP release, motif **IVa** advances toward P_n+4_ while motif **V** re-approaches P_n+2_. This coordinated progression suggests that motif **IVa** steps from P_n_ to P_n+2_ by traversing P_n+1_ with assistance from motif **V**. The observed alternation between stable attachment and diffusive sampling, particularly in motif **IVa**, aligns with a small-step inchworm mechanism. Similar translocation behavior has been proposed in other *Orthoflavivirus* helicases,^16,20,38^ although a Brownian ratchet model has also been suggested for ZIKV NS3 helicase.^18^

ATP-dependent coupling between motif **VIL** and RNA-cleft contacts is mediated through specific phosphate interactions that shift over the hydrolysis cycle. To pinpoint the RNA-phosphate interactions most associated with motif **VIL**’s nucleotide affinity, we performed residue-level correlation analysis across hydrolysis states (Figure 5A). We further decomposed these correlations by examining interactions with the two primary ADP-contacting residues of motif **VIL**, R461 (Figure 5B) and R464 (Figure 5C). Among all contacts, the RNA phosphate affinity of D410 (motif **V**) and K389 (motif **IVa**) show only weak correlation with motif **VIL**’s ADP affinity (Figure 5A). For R388 (motif **IVa**), the correlation coefficients between its P_1_ affinity and motif **VIL** ADP affinity are –0.19, 0.06, and –0.31 in the ssRNA+ATP, ssRNA+ADP+P_i_, and ssRNA+ADP states, respectively. This behavior is primarily driven by coupling to R461 (Figure 5B), suggesting an allosteric link between R388 and R461. For R388’s interaction with P_3_, the correlation coefficients were 0.19, 0.27, and –0.28 across the same states, influenced by both R461 and R464 (Figure 5B–C). In contrast, the P_2_ affinity of T409 (motif **V**) exhibited the strongest and most consistent correlation with motif **VIL**’s ADP affinity, increasing in magnitude along the hydrolysis cycle. The correlation coefficients are –0.10 (ssRNA+ATP), –0.33 (ssRNA+ADP+P_i_), and –0.43 (ssRNA+ADP), reflecting increasingly tight inverse coupling. This trend is primarily mediated by R464 in the ssRNA+ADP+P_i_ state and by R461 in the ssRNA+ADP state, underscoring the functional significance of T409(**V**)–**VIL** coupling.

**Figure 5:**
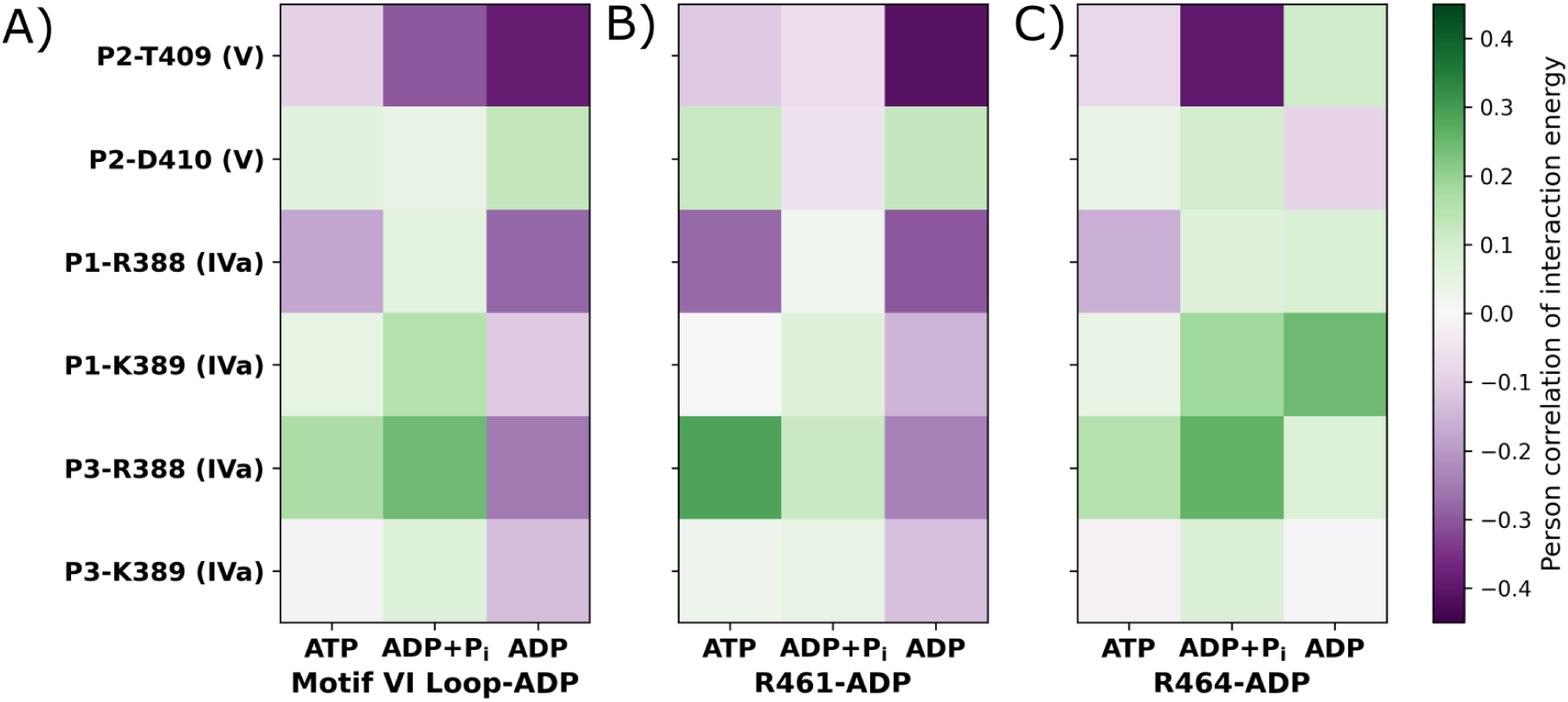
Correlation between RNA phosphate affinity and ADP interaction energy of motif VIL residues. Key RNA-binding residues were identified from ATP-dependent changes in RNA phosphate affinity (see Figure 3). The selected phosphate-residue pairs include: P_2_–T409 (motif **V**), P_2_–D410 (motif **V**), P_1_–R388 (motif **IVa**), P_1_–K389 (motif **IVa**), P_3_–R388 (motif **IVa**), and P_3_–K389 (motif **IVa**). (A) Correlation coefficients between the RNA phosphate interaction energies of these residue pairs and the ADP interaction energy of motif **VIL**, computed for the ssRNA+ATP, ssRNA+ADP+P_i_, and ssRNA+ADP states. (B, C) Correlations between RNA phosphate affinities of the same residues and the ADP interaction energies of R461 and R464, respectively—two residues previously shown to dominate the ADP binding of motif **VIL**.^19^

Disruption of the coupling between motif **VIL** and RNA phosphate affinity in motif **V** impairs the conformational transitions required for RNA translocation. In simulations of motif **VIL** loop mutant, we observed no ATP-dependent changes in phosphate affinity at the key RNA-cleft contacts of motifs **V** and **IVa** (Figure S5), in contrast to the behavior of the wild-type enzyme. This absence of phosphate affinity modulation coincides with the loss of correlation between the P_2_ affinity of T409 (motif **V**) and the ADP affinity of motif **VIL** (Figure S6). In the wild-type enzyme, this correlation supports a functional pathway for signal transmission from the ATP-pocket to the RNA-cleft. Based on these observations, we propose that motif **V** serves as both a receiver of nucleotide-affinity changes from motif **VIL** and a responder that modulates RNA phosphate affinity to enable translocation. These findings underscore the central role of motif **VIL**–**V** coupling in enabling efficient, directional RNA translocation through hydrolysis-state-dependent phosphate affinity shifts.

### Allosteric correlation between motif VI and RNA affinity of motif V

ATP-dependent structural coupling between motif **VIL** and motif **V** mirrors their functional coordination in substrate binding. To evaluate whether changes in RNA and nucleotide affinity are accompanied by coordinated structural motion, we calculated the positional correlation between the centers of mass of backbone and sidechain atoms in motifs **VIL** and **V** (Figure 6). In the ssRNA state, correlation coefficients range from –0.22 to 0.60 for backbone atoms (Figure 6A) and –0.37 to 0.52 for sidechain atoms (Figure 6E), indicating moderate coupling. These correlations weaken in the ssRNA+ATP state, with backbone coefficients between –0.07 and 0.44 (Figure 6B) and sidechain correlations from –0.33 to 0.41 (Figure 6F). In the ssRNA+ADP+P_i_ state, the correlation partially recovers, with backbone values ranging from –0.30 to 0.52 (Figure 6C) and sidechain correlations from –0.33 to 0.59 (Figure 6G). Finally, in the ssRNA+ADP state, the correlation strengthens further, especially for the backbone (–0.16 to 0.60; Figure 6D), while sidechain values range from –0.39 to 0.46 (Figure 6H). These results indicate that structural coordination between motifs **VIL** and **V** is dynamically modulated by the hydrolysis state and parallels the trends observed in their substrate binding affinity.

**Figure 6:**
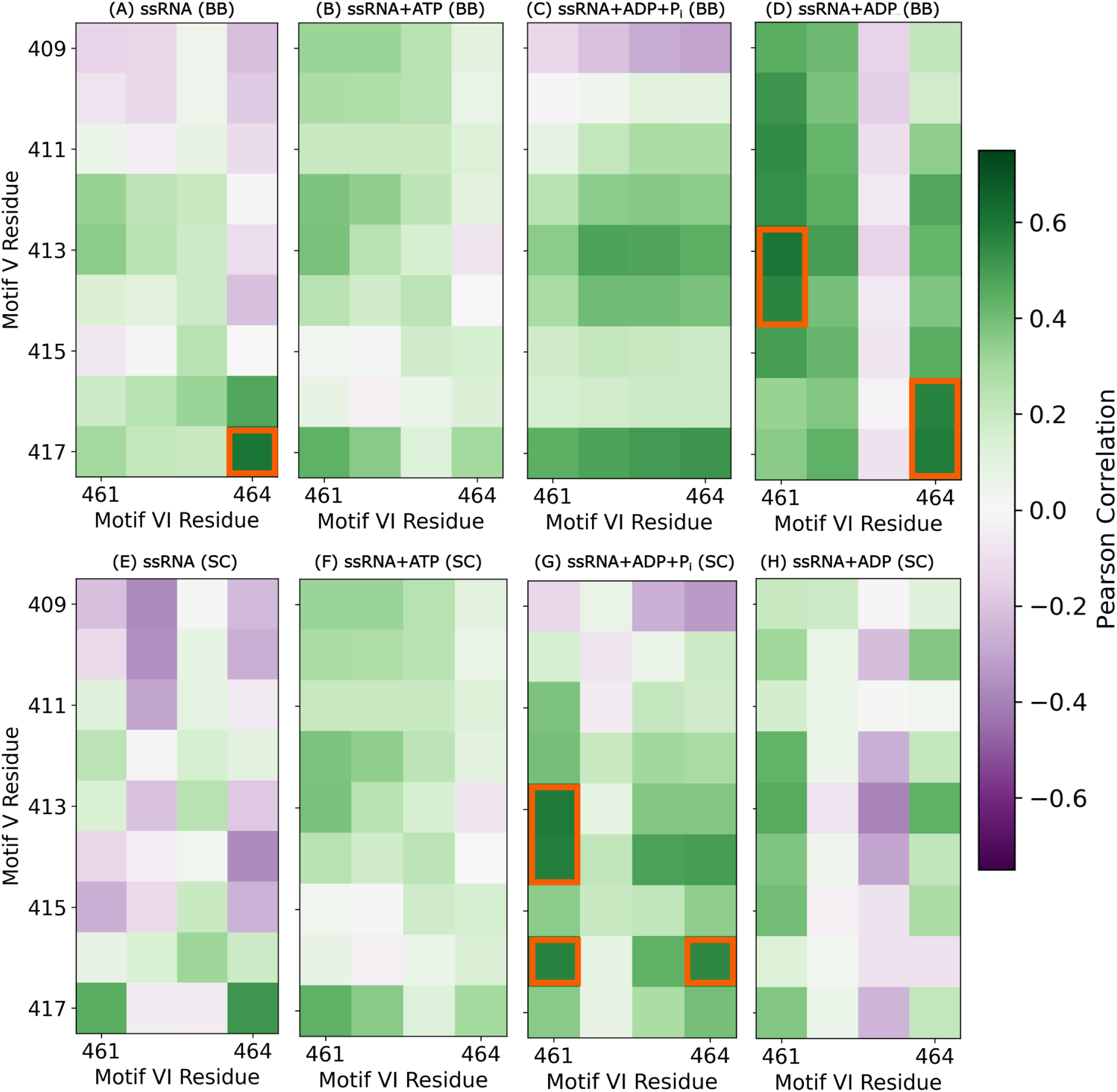
Identification of key correlated residue pairs between motif V and motif VIL via positional correlation analysis. Positional correlations of the centers of mass for backbone (BB; top row) and sidechain (SC; bottom row) atoms were computed after global alignment of each structure within its respective hydrolysis state ensemble. Panels represent: (A, E) ssRNA, (B, F) ssRNA+ATP, (C, G) ssRNA+ADP+P_i_, and (D, H) ssRNA+ADP states. Orange squares highlight residue pairs with correlation coefficients greater than 0.55 or less than –0.55, indicating strong positive or negative correlations, respectively.

R461–E413 interaction serves as a key structural link between motifs **VIL** and **V** during ATP hydrolysis. In our structural correlation analysis, residue pairs exhibiting correlation coefficients greater than |0.55| were marked as strongly coupled and are highlighted in orange in Figure 6. Among these, the R461–E413 and R464–A416 pairs are particularly notable, with strong correlations emerging in the ssRNA+ADP+P_i_ and ssRNA+ADP states. Notably, the nature of these correlations shifts from sidechain–sidechain to backbone–backbone interactions between states. Structural proximity between R461 and E413 (compared to R464–A416) suggests R461–E413 as a more plausible site of direct interaction (Figure S7). To further probe this interaction, we quantified hydrogen bond (H-bond) occupancy between the R461 sidechain (SC) and both the backbone (BB) and sidechain (SC) of E413. No significant H-bonding was observed in the ssRNA or ssRNA+ATP states (Figure S8). In the ssRNA+ADP+P_i_ state, R461(SC) formed an H-bond with E413(BB) in 72.10 ± 4.78% of frames, while no bonding with E413(SC) is detected (Figure 7A). Upon transition to the ss-RNA+ADP state, this interaction shifted: R461(SC)–E413(BB) H-bond occupancy dropped to 6.06 ± 3.95%, whereas R461(SC)–E413(SC) H-bonding increased to 54.94 ± 14.73% (Figure 7B). This switching of interaction partners coincides with enhanced ADP affinity of R461 observed in this transition.^19^ Importantly, this conformational switch is absent in simulations of the motif **VIL** mutant (Figure S9), further supporting its functional relevance.

**Figure 7:**
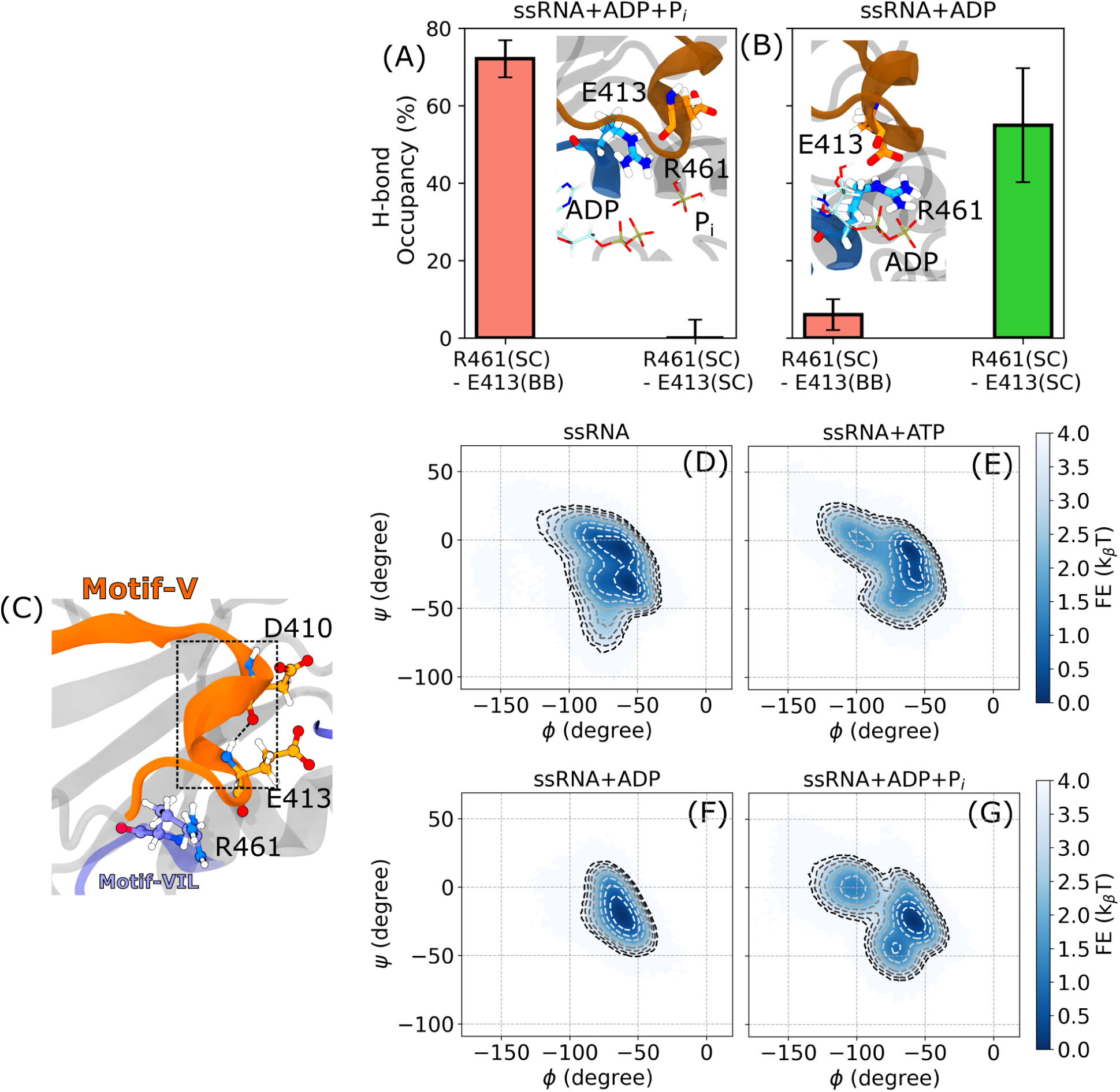
Correlation between R461 and E413 modulates ATP-dependent structural sampling of the motif V 3_10_ helix. (A, B) Hydrogen bond (H-bond) sampling between the sidechain of R461 (acceptor) and either the backbone or sidechain of E413 (donor) in the ssRNA+ADP+P_i_ (A) and ssRNA+ADP (B) states. No H-bond formation is observed between these residues in the ssRNA or ssRNA+ATP states (Figure S8). Insets show structural snapshots illustrating the relative positions of E413 and R461. (C) A representative structure highlighting the motif **V** 3_10_ helix, spanning residues D410 to E413. (D–G) Two-dimensional free energy surfaces projected along the *ϕ* and *ψ* dihedral angles of E413 for the (D) ssRNA, (E) ssRNA+ATP, (F) ssRNA+ADP, and (G) ssRNA+ADP+P_i_ states. Contours are drawn at intervals of 0.5 k_B_T, from 0 to 4 k_B_T. ATP-dependent shifts in sampling indicate structural heterogeneity in the 3_10_ helix regulated by the R461–E413 interaction.

ATP-dependent interaction between E413 (motif **V**) and R461 (motif **VIL**) influences the structural heterogeneity of the 3_10_ helix in motif **V**. Residue E413 resides within this helix, which also includes D410 and its preceding residue T409—both of which interact with ssRNA (Figure 7C). To evaluate how the E413–R461 interaction affects the 3_10_ helix conformation, we computed two-dimensional free energy surfaces in the *ϕ*–*ψ* dihedral space (Figure 7D-G). In the ssRNA state, the 3_10_ helix samples two minima located at (−55^◦^, −33.5^◦^) and (−61^◦^, −6.5^◦^), indicating a broader distribution in *ϕ* angles (Figure 7D). The latter minimum is also sampled in the ssRNA+ATP state (Figure 7E), though a distinct intermediate at (−95^◦^, 0^◦^) replaces the second minimum observed in the ssRNA state. In the ssRNA+ADP+P_i_ state, the helix adopts a new minimum at (−59^◦^, −24.5^◦^), along with two additional intermediates at (−71^◦^, −44.3^◦^) and (−103^◦^, 0^◦^) (Figure 7G), suggesting a structural shift linked to R461–E413 interaction. Following P_i_ release, in the ssRNA+ADP state—where the R461–E413 interaction site shifts—the helix samples only a single minimum at (−61^◦^, −20.9^◦^) (Figure 7F). In motif **VIL** mutants, which lack this ATP-dependent switching of the R461–E413 interaction, no change in 3_10_ helix sampling is observed across hydrolysis states (Figure S10).

## Conclusions

This study investigated the ATP-dependent coupling mechanism that links nucleotide affinity in the ATP-binding pocket to single-stranded RNA (ssRNA) affinity in the RNA-cleft of the West Nile Virus (WNV) NS3 helicase (NS3h). By elucidating the structural basis of this coupling, we aimed to improve our understanding of the ATP hydrolysis-driven translocation process—an essential step in viral genome replication. Using previously generated simulation data for WNV NS3h bound to ssRNA in distinct ATP hydrolysis states (ssRNA, ssRNA+ATP, ssRNA+ADP+P_i_, and ssRNA+ADP),^19^ we built upon earlier findings that identified the motif **VI** loop as a nucleotide valve controlling ADP affinity across the catalytic cycle. To assess whether this coupling mechanism is functionally conserved, we further analyzed motif **VI** loop mutants (D471E, D471L), which are known to disrupt nucleotide binding and attenuate viral replication.

Our correlation analyses revealed that both motif **I** and the motif **VI** loop exhibit coupling between ADP affinity and ssRNA phosphate affinity. However, the motif **VI** loop uniquely displays an ATP-dependent shift in this correlation, particularly with phosphate interactions involving motif **IVa** and **V**. At the residue level, R388 and K389 of motif **IVa**, along with T409 and D410 of motif **V**, undergo a sequential change in ssRNA phosphate affinity across the hydrolysis cycle, consistent with the translocation of one phosphate unit per cycle. Notably, the ssRNA affinity of motif **V** is strongly correlated with the ADP affinity of R461 and R464 in motif **VI**, previously identified as arginine fingers essential for NTPase activity and nucleotide gating.

Further structural analysis revealed a hydrogen bond–mediated connection between E413 (motif **V**) and R461 (motif **VI**), forming an allosteric bridge that links ATP-pocket dynamics to the RNA-cleft. This interaction modulates the structural heterogeneity of the 3_10_-helix in motif **V**, a conformational feature directly involved in ssRNA binding. Importantly, this E413–R461 interaction and its downstream structural effects are absent in the motif **VI** loop mutants, reinforcing its functional significance.

Taken together, our findings define a mechanistic conduit by which ATP-dependent conformational changes originating at motif **VI** propagate through E413 to reshape motif **V** and modulate ssRNA affinity. These insights not only advance the molecular understanding of viral RNA helicase function but also identify a structurally and energetically conserved allosteric pathway that could serve as a promising target for antiviral drug development against WNV and related *Orthoflaviviruses*.

## Acknowledgement

Research reported in this manuscript was supported by the National Institute for Allergic and Infectious Diseases of the National Institute of Health under award number R01AI166050. Computational resources for this project were provided by: (1) the High Performance Computing Center at Oklahoma State University supported in part through the National Science Foundation Grant OAC-1531128 and (2) Purdue Anvil under ACCESS project number BIO220160.

## Notes

### Competing Interest Statement

The authors have declared no competing interest.

