## Supporting Information for "Allosteric Regulation of RNA Affinity by Motif V-VI Coupling in West Nile Virus NS3 Helicase"

Priti Roy and Martin McCullagh*

*Department of Chemistry, Oklahoma State University, Stillwater, OK, 74074, USA*

**Table S1: Occurrence probability of conserved RNA contacts in the ssRNA state of WNV NS3h mutants. Wild-type data is included for comparison. A 5 Å of cut-off distance is used between atoms of protein and ssRNA.**

| Motif | Residue | Wild-type | D471E | D471L |
|---|---|---|---|---|
| **Ia** | P224 | 99% | 97.73% | 100% |
| **Ia** | T225 | 99% | 98.18% | 100% |
| **Ia** | R226 | 99% | 99.99% | 100% |
| **II** | T290 | 64% | 0.15% | 68.27% |
| **II** | D291 | 99% | 59.93% | 97.98% |
| **IV** | S365 | 35% | 99.85% | 97.51% |
| **IVa** | R388 | 99% | 99.96% | 100% |
| **V** | D410 | 97% | 7.64% | 100% |

**Table S2: Occurrence probability of conserved RNA contacts in the ssRNA+ATP state of WNV NS3h mutants. Wild-type data is included for comparison. A 5 Å of cut-off distance is used between atoms of protein and ssRNA.**

| Motif | Residue | Wild-type | D471E | D471L |
|---|---|---|---|---|
| **Ia** | P224 | 99% | 99.76% | 98.47% |
| **Ia** | T225 | 81% | 99.65% | 99.86% |
| **Ia** | R226 | 99% | 100% | 100% |
| **II** | T290 | 8% | 14.14% | 23%% |
| **II** | D291 | 99% | 99.27% | 94.05% |
| **IV** | P364 | 93% | 80.03% | 99.52% |
| **IV** | S365 | 99% | 75.84% | 97.81% |
| **IV** | V366 | 98% | 73.35% | 96% |
| **IV** | K367 | 56% | 72.53% | 40.6% |
| **IVa** | S387 | 99% | 1.31% | 96.79% |
| **IVa** | R388 | 99% | 99.99% | 100% |
| **IVa** | K389 | 63% | 14.12% | 32.19% |
| **V** | T409 | 99% | 88.62% | 100% |
| **V** | D410 | 99% | 27.62% | 99.98% |
| **V** | I411 | 99% | 99.58% | 100% |

**Table S3: Occurrence probability of conserved RNA contacts in the ssRNA+ADP+$P_i$ state of WNV NS3h mutants. Wild-type data is included for comparison. A 5 Å of cut-off distance is used between atoms of protein and ssRNA.**

| Motif | Residue | Wild-type | D471E | D471L |
|---|---|---|---|---|
| **Ia** | P224 | 99% | 69.91% | 99.03% |
| **Ia** | T225 | 99% | 91.47% | 99.94% |
| **Ia** | R226 | 99% | 100% | 100% |
| **II** | D291 | 99% | 80.28% | 99.75% |
| **IV** | S365 | 97% | 99.69% | 36.72% |
| **IV** | V366 | 99% | 97.4% | 99.24% |
| **IV** | K367 | 84% | 14.07% | 4.95% |
| **IVa** | S387 | 99% | 99.83% | 99.67% |
| **IVa** | R388 | 99% | 100% | 99.88% |
| **IVa** | K389 | 93% | 11.81% | 39.04% |
| **V** | T409 | 99% | 100% | 99.99% |
| **V** | D410 | 99% | 57.53% | 85.74% |
| **V** | I411 | 99% | 100% | 100% |

**Table S4: Occurrence probability of conserved RNA contacts in the ssRNA+ADP state of WNV NS3h mutants. Wild-type data is included for comparison. A 5 Å of cut-off distance is used between atoms of protein and ssRNA.**

| Motif | Residue | Wild-type | D471E | D471L |
|---|---|---|---|---|
| **Ia** | P224 | 99% | 100% | 100% |
| **Ia** | T225 | 99% | 100% | 100% |
| **Ia** | R226 | 99% | 100% | 100% |
| **II** | T290 | 63% | 65.40% | 1.12% |
| **II** | D291 | 99% | 99.68 | 92.65% |
| **IV** | P364 | 99% | 100% | 76.63% |
| **IV** | S365 | 99% | 98.96% | 23.83% |
| **IV** | V366 | 99% | 98.02% | 24.64% |
| **IV** | K367 | 95% | 62.02% | 7.92% |
| **IVa** | R388 | 99% | 99.99% | 100% |
| **V** | T409 | 98% | 100% | 100% |
| **V** | I411 | 98% | 99.99% | 100% |

**Table S5: Occurrence probability of conserved ATP contacts in the ssRNA+ATP state of WNV NS3h mutants. Wild-type data is included for compariso. A 5 Å of cut-off distance is used between atoms of protein and ATP.**

| **Motif** | **Residue** | **Wild-type** | **D471E** | **D471L** |
|---|---|---|---|---|
| **I** | H195 | 61% | 0.25% | 100% |
| **I** | P196 | 99% | 22.55% | 100% |
| **I** | G197 | 99% | 99.75% | 100% |
| **I** | A198 | 99% | 100% | 100% |
| **I** | G199 | 99% | 100% | 100% |
| **I** | K200 | 99% | 100% | 100% |
| **I** | T201 | 99% | 100% | 100% |
| **V** | N417 | 92% | 97.45% | 70.73% |
| **VI** | R459 | 99% | 0.01% | – |
| **VI** | R462 | 99% | 21.58% | – |

**Table S6: Occurrence probability of conserved ADP+$P_i$ contacts in the ssRNA+ADP+$P_i$ state of WNV NS3h mutants. Wild-type data is included for compariso. A 5 Å of cut-off distance is used between atoms of protein and ADP+$P_i$.**

| **Motif** | **Residue** | **Wild-type** | **D471E** | **D471L** |
|---|---|---|---|---|
| **I** | H195 | 86% | 99.9% | 1.82% |
| **I** | P196 | 99% | 99.85% | 75.26% |
| **I** | G197 | 99% | 100% | 33.40% |
| **I** | A198 | 99% | 100% | 18.48% |
| **I** | G199 | 99% | 100% | 92.80% |
| **I** | K200 | 99% | 100% | 90.54% |
| **I** | T201 | 99% | 100% | 100% |
| **II** | E286 | 98% | 96.55% | – |
| **V** | G415 | 99% | 0.11% | 50.47% |
| **V** | N417 | 90% | 32.13% | 97.9% |
| **VI** | Q455 | 98% | – | – |
| **VI** | R459 | 99% | – | – |
| **VI** | R462 | 99% | 0.18% | 35.66% |
| **VI** | N465 | 99% | 2.42% | 34.09% |

**Table S7: Occurrence probability of conserved ADP contacts in the ss-RNA+ADP state of WNV NS3h mutants. Wild-type data is included for compariso. A 5 Å of cut-off distance is used between atoms of protein and ADP.**

| Motif | Residue | Wild-type | D471E | D471L |
|---|---|---|---|---|
| **I** | P196 | 97% | 86.66% | 99.85% |
| **I** | G197 | 99% | 91.77% | 100% |
| **I** | A198 | 99% | 99.96% | 100% |
| **I** | G199 | 99% | 100% | 100% |
| **I** | K200 | 99% | 100% | 100% |
| **I** | T201 | 99% | 100% | 100% |
| **II** | E286 | 68% | 5.83% | 1.28% |
| **V** | G415 | 23% | 78.21% | 1.64% |
| **V** | N417 | 94% | 77.42% | 74.88% |
| **VI** | R459 | 97% | 0.54% | 0.59% |
| **VI** | R461 | 70% | 0.02% | 0.37 |
| **VI** | N465 | 97% | 9.7% | 77.38% |

**Table S8: Probability of oxygen atoms participated in the octahedral coordination of $Mg^{2+}$ in WNV NS3H mutant. This calculation is performed for all hydrolysis states of D471L. The octahedral shell is defined by oxygen atoms with 2.5 Å of $Mg^{2+}$.**

| **Atom Name** | **ssRNA+ATP** | **ssRNA+ADP+$P_i$** | **ssRNA+ADP** |
|---|---|---|---|
| $OG1_{T200}$ | 99.97% | 99.63% | 99.84% |
| $O2G_{ATP}$ | 100% | – | – |
| $O1B_{ATP}$ | 100% | – | – |
| $O1B_{ADP}$ | – | 100% | 100% |
| $O3_{P_i}$ | – | 100% | – |
| $O_{W1}$ | 100% | 100% | 100% |
| $O_{W2}$ | 100% | 100% | 100% |
| $O_{W3}$ | 100% | 100% | 100% |
| $O_{W4}$ | – | – | 100% |

Figure S1: **2D histogram of backbone dihedral angles of the entire NS3h** The $\phi$ and $\psi$ are denoted as backbone dihedrals and these values are calculated for each hydrolysis state of Wild-type and NS3h mutants. Units in degrees.

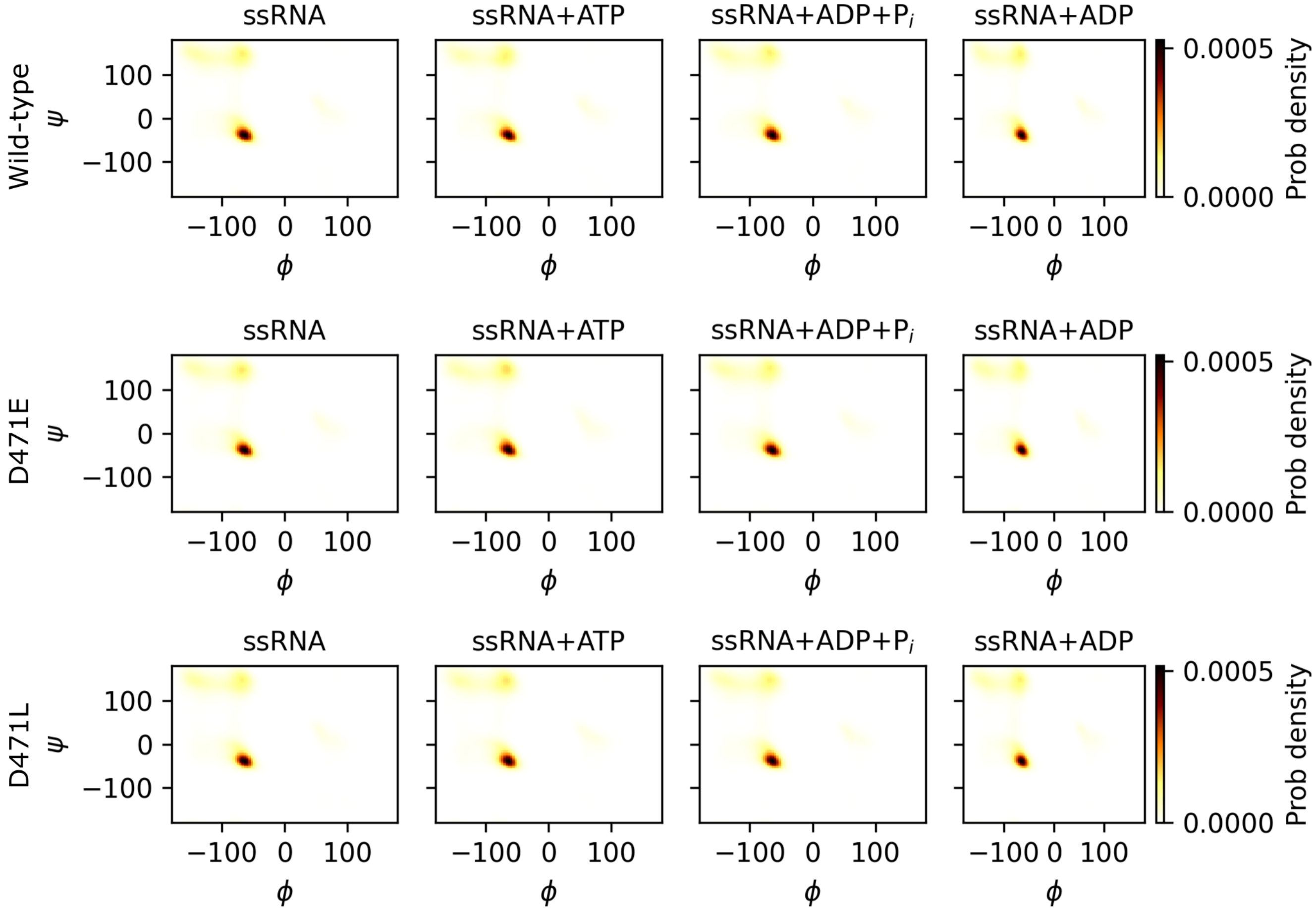

Figure S2: **ATP-dependent correlation between nucleotide and RNA affinity in WNV NS3h D471E.** (A–C) Residue-wise cross-correlation plots comparing nucleotide (ADP) and RNA(P) binding affinities for three hydrolysis states: (A) ssRNA+ATP (pre-hydrolysis), (B) ssRNA+ADP+$P_i$ (post-hydrolysis I), and (C) ssRNA+ADP (post-hydrolysis II). The black rectangle in each plot denote significant correlation region observed in Wild-type. For analysis details see main text.

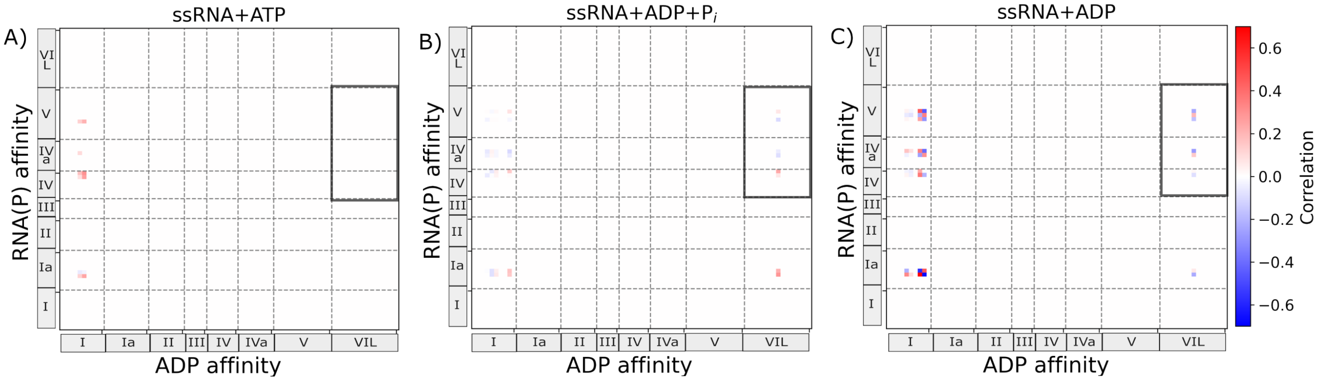

Figure S3: **ATP-dependent correlation between nucleotide and RNA affinity in WNV NS3h D471L.** (A–C) Residue-wise cross-correlation plots comparing nucleotide (ADP) and RNA(P) binding affinities for three hydrolysis states: (A) ssRNA+ATP (pre-hydrolysis), (B) ssRNA+ADP+$P_i$ (post-hydrolysis I), and (C) ssRNA+ADP (post-hydrolysis II). The black rectangle in each plot denote significant correlation region observed in Wild-type. For analysis details see main text.

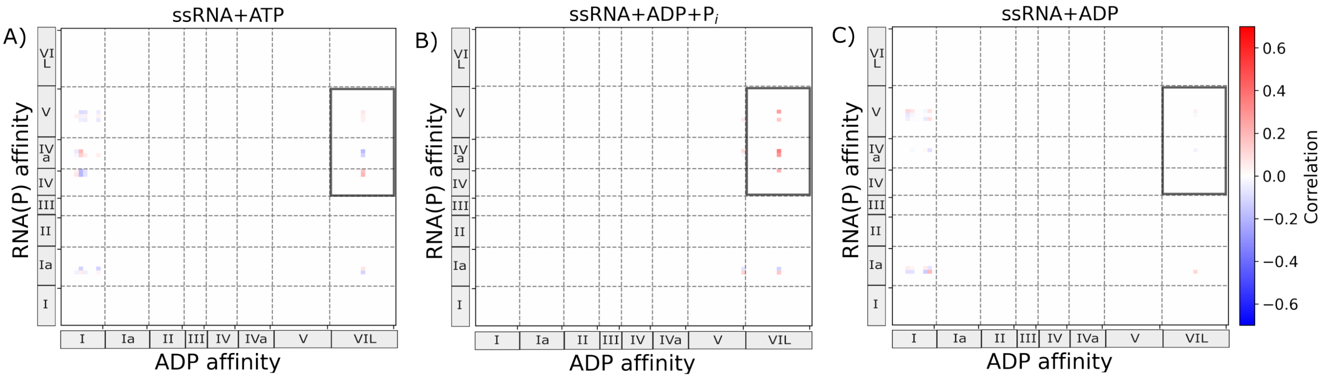

Figure S4: **Residue level phosphate affinity change at RNA-cleft motifs Ia, IV, IVa and V in Wild-type**. The binding affinity change for each protein residue for phosphates $P_1$ to $P_6$ are presented from top to bottom panel. The deviation correspond to difference of the ensemble averages, those are computed from entire NPT ensemble of the respective system. The errorbar represents propagated standard deviation of two compared systems where standard deviations are computed from the averages of 50k frames chunks. See main text for details. The units in kcal.mol$^{-1}$

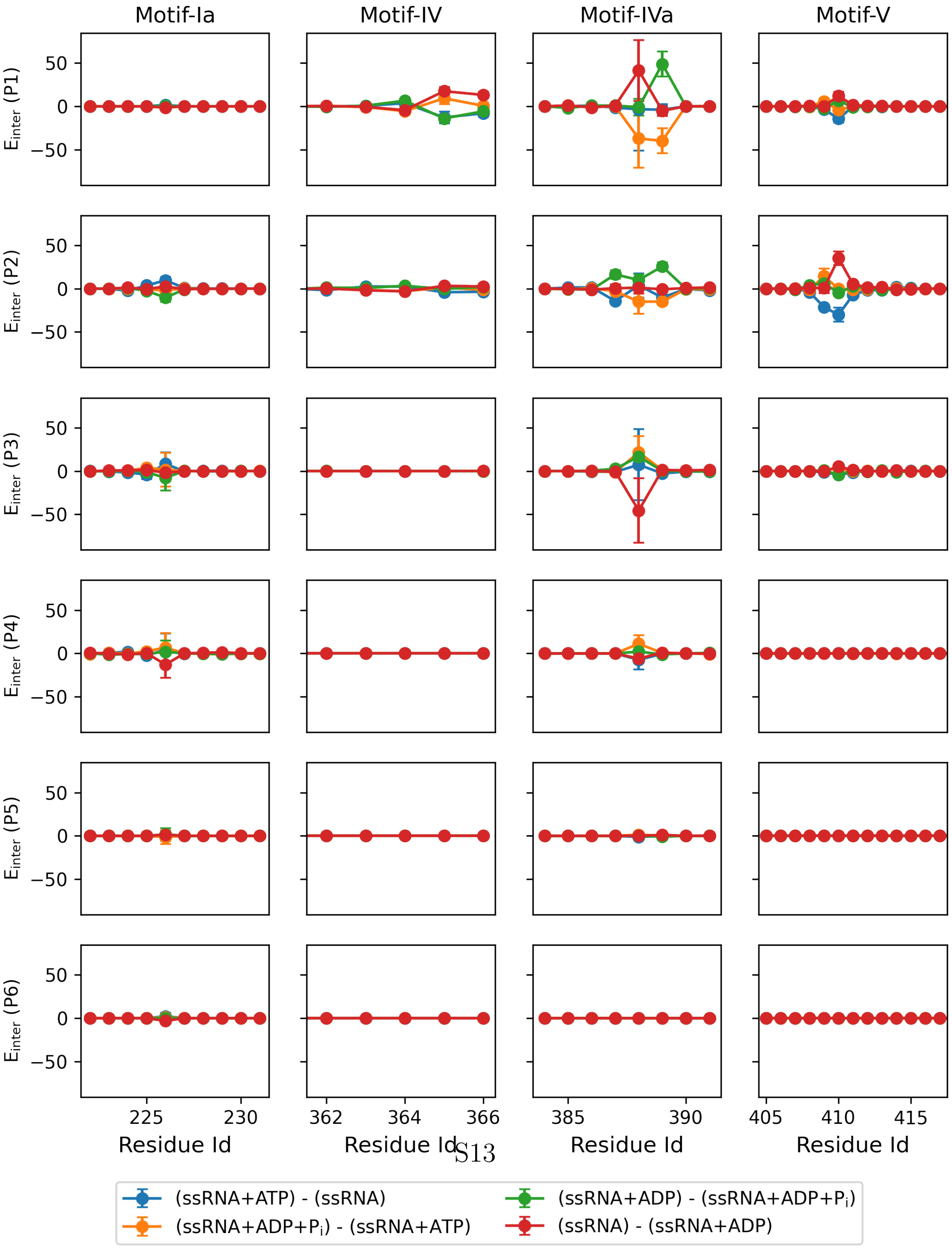

Figure S5: **ATP-dependent ssRNA phosphate affinity change for motif IVa and V in the D471L**. Residue wise differential binding enthalpy ($\Delta E_{inter}$) of motif **V** for **P2**, motif **IVa** for **P1** and motif **IVa** for **P3** shown in left, middle and right columns respectively. The deviation corresponds to difference of the averages, those are computed from entire NPT ensemble of the respective states. The error bar represents propagated standard deviation of two compared states where standard deviations are computed from the averages of 50k frames chunks. (A-C) These plots mark the change in affinity while protein transition from ssRNA to ssRNA+ATP state. (D-E) Changes in RNA phosphate affinity after protein moving from ssRNA+ATP state to ssRNA+ADP+$P_i$ state. (G-I) RNA phosphate affinity change due to transition of protein from ssRNA+ADP+$P_i$ to ssRNA+ADP state. (J-L) Changes in RNA phosphate affinity while protein move from ssRNA+ADP state of previous cycle to the ssRNA state of next cycle. Units of $\Delta E_{inter}$ is in kcal·mol$^{-1}$

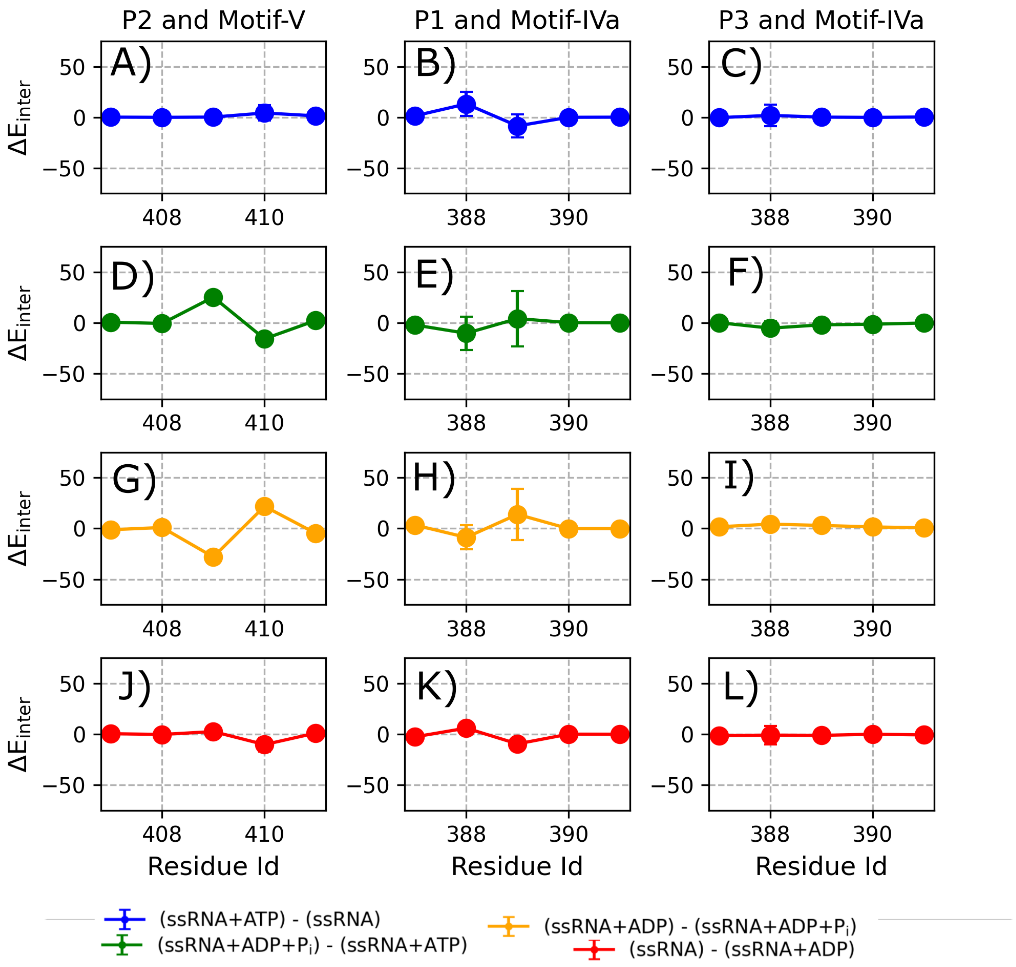

Figure S6: **Correlation of ADP affinity of motif VIL and key wild-type RNA contacts in D471L** The RNA interacting key residues were identified from ATP-dependent changes in the protein residue and RNA phosphates affinity (Figure **??**) in the wild-type. The key interacting residue pairs are: P2 and T409 of motif **V**, P2-D410 of motif **V**, P1 and R388 from motif **IVa**, P1 and K389 from motif **IVa**, P3 and R388 from motif **IVa**, P3 and K389 from motif **IVa**. Correlation between the interaction energy of these pairs in the RNA-cleft and ADP interaction energy of motif **VIL** calculated for hydrolysis states ATP, ADP+$P_i$ and ADP.

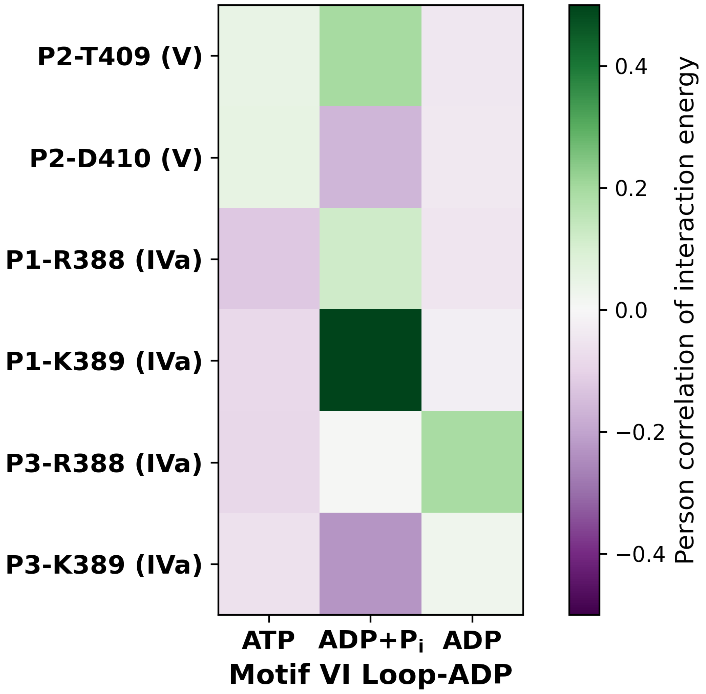

Figure S7: **Time evolution of distances of residue pairs between motif V and VIL**. The plots represent the center-of-mass distances for backbones or sidechains for residue pairs showed correlation coefficient equal or greater than 0.55 in the hydrolysis states (see main text). These time traces demonstrate the stability or fluctuation along with spatial proximity of the motif V and VI residue pairs. The distance units in Å.

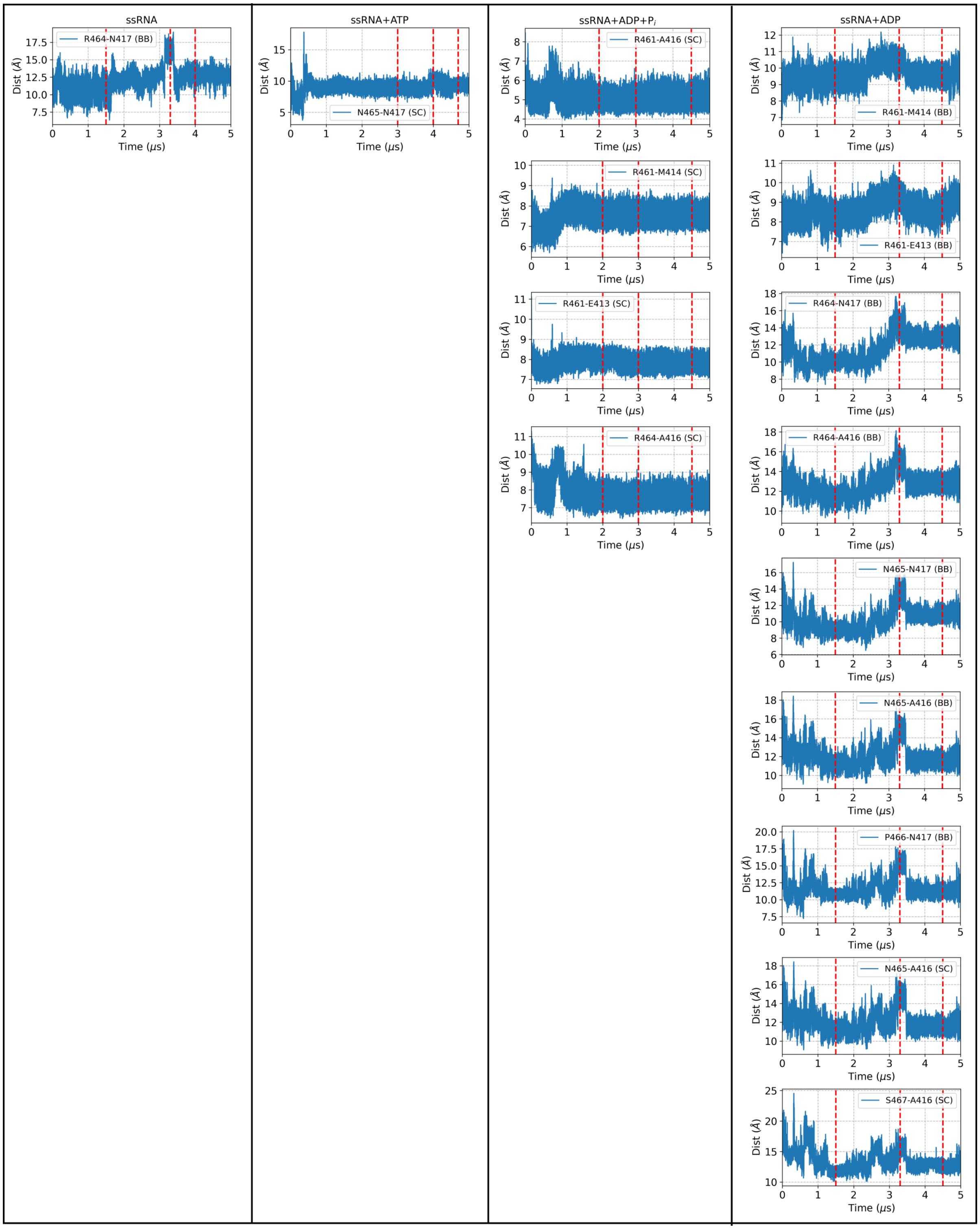

Figure S8: **Sampling of E413-R461 hydrogen bond in the Wild-type ssRNA and ssRNA+ATP states.** The backbone and sidechain atoms of E413 act as an acceptor while the sidechain atoms of R461 act as a donor.

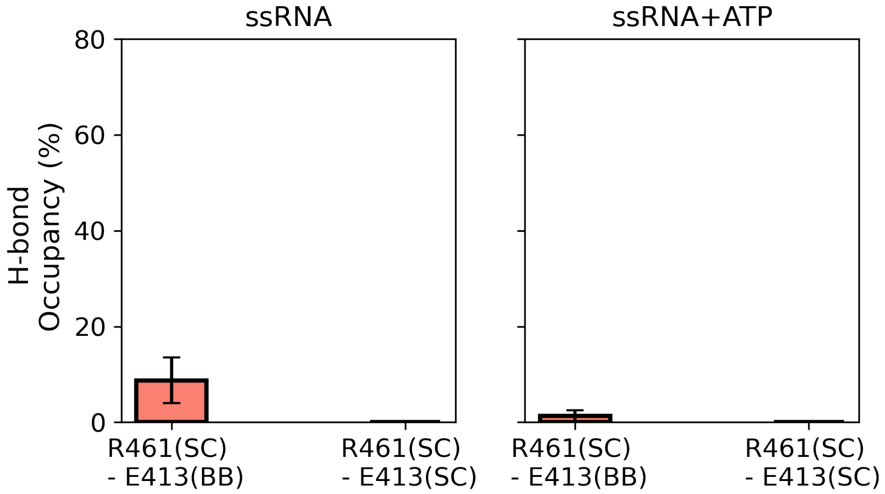

Figure S9: **Sampling of E413-R461 hydrogen bond in the hydrolysis states of D471L.** The backbone and sidechain atoms of E413 act as an acceptor while the sidechain atoms of R461 act as a donor.

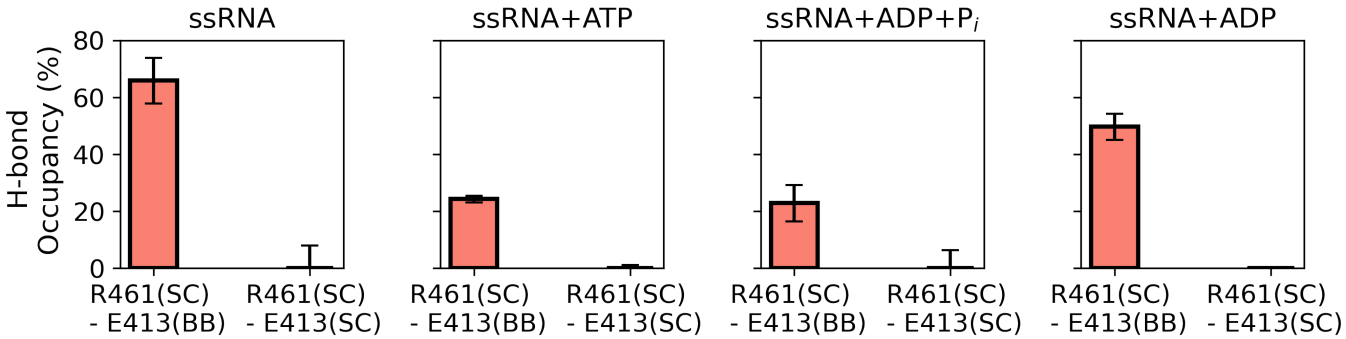

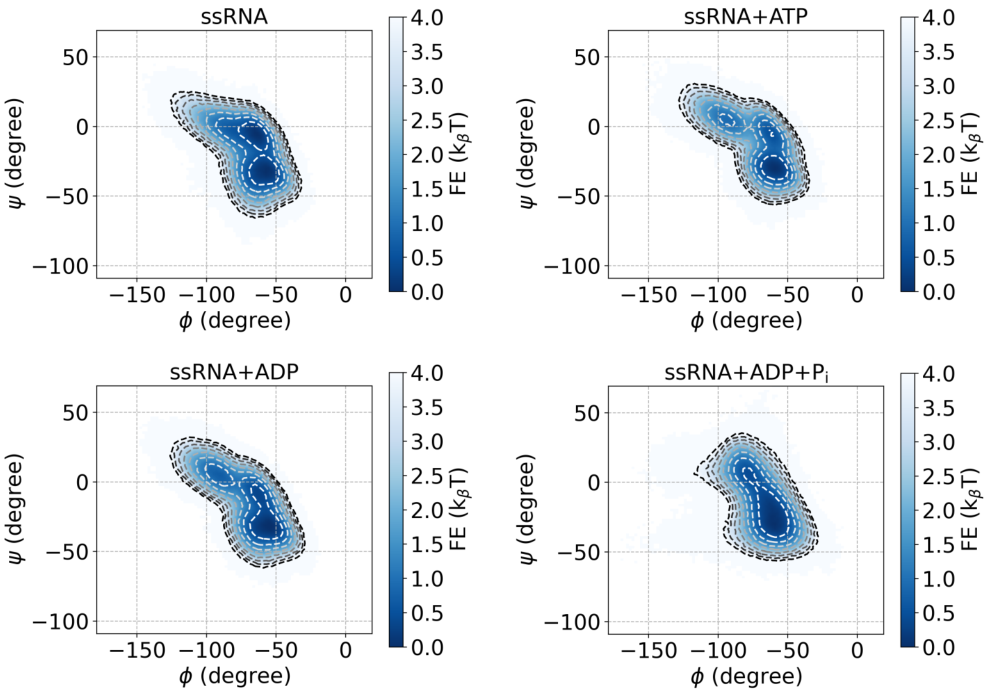


Figure S10: **Structural sampling of $3_{10}$-helix of motif V in the hydrolysis states of D471L.** These 2D free energy plots show ATP-dependent sampling of $\phi$ and $\psi$ angles of $3_{10}$-helix. Contour lines are plotted with an interval of 0.5 $k_\beta$T, ranging from 0 to 4 $k_\beta$T.
